# Tropomyosin 3.5 protects F-actin networks required for tissue biomechanical properties

**DOI:** 10.1101/357509

**Authors:** Catherine Cheng, Roberta B. Nowak, Michael B. Amadeo, Sondip K. Biswas, Woo-Kuen Lo, Velia M. Fowler

**Author notes:** Corresponding Author: Velia M. Fowler Department of Molecular Medicine The Scripps Research Institute 10550 N. Torrey Pines Road, MB114 La Jolla, CA 92037.

## Abstract

Tropomyosins (Tpms) stabilize F-actin and regulate interactions with other actin-binding proteins. The eye lens changes shape in order to fine focus light to transmit a clear image, and thus lens organ function is tied to its biomechanical properties, presenting an opportunity to study Tpm functions in tissue mechanics. The major mouse lens Tpm is Tpm3.5 (TM5NM5), a previously unstudied isoform. Decreased levels of Tpm3.5 lead to softer and less mechanically resilient lenses that are unable to resume their original shape after compression. While cell organization and morphology appear unaffected, Tmod1 dissociates from the membrane in Tpm3.5-deficient lens fiber cells resulting in reorganization of the spectrin-F-actin and α-actinin-F-actin networks at the membrane. These rearranged F-actin networks appear to be less able to support mechanical load and resilience leading to an overall change in tissue mechanical properties. This is the first *in vivo* evidence that Tpm is essential for cell biomechanical stability in a load-bearing non-muscle tissue and indicates that Tpm3.5 protects mechanically stable, load-bearing F-actin *in vivo*.

**Summary:** Tropomyosin 3.5 stabilizes F-actin in eye lens fiber cells and promotes normal tissue biomechanical properties. Tpm3.5 deficiency leads to F-actin network rearrangements and decreased lens stiffness and resilience.

## Introduction

Tropomyosins (Tpms) are conserved F-actin binding proteins that stabilize filaments and regulate their interactions with a variety of actin-binding proteins, including cofilin/ADF filament severing proteins, α-actinin and fimbrin (plastin) filament crosslinking proteins, tropomodulin (Tmod) filament pointed-end-capping proteins, and force-producing myosin motors (Christensen et al., 2017; Gunning et al., 2015; Hitchcock-DeGregori and Barua, 2017; Kostyukova, 2008; Nakano and Mabuchi, 2006; Ono and Ono, 2002; Winkelman et al., 2016; Yamashiro et al., 2012). Tpms are expressed in all animals and fungi (Barua et al., 2011; Cranz-Mileva et al., 2013), and in mammals, Tpms are expressed from 4 genes with alternative splicing that produces more than 40 variants in different tissues (Geeves et al., 2015; Pittenger et al., 1994; Vindin and Gunning, 2013). *In vitro* studies have demonstrated that F-actin assembly, elongation and disassembly rates depend on Tpm (Gunning et al., 2015; Hitchcock-DeGregori and Barua, 2017) and that Tpm isoforms direct the assembly of different types of F-actin populations (Gateva et al., 2017; Janco et al., 2016). Tpm1.6 (Tm2) and Tpm1.7 (Tm3) protect F-actin from ADF/cofilin-mediated disassembly, while Tpm3.1 (TM5NM1), Tpm3.2 (TM5NM2), and Tpm4.2 (Tm4) stimulate myosin IIA binding to F-actin but do not protect filaments from disassembly (Gateva et al., 2017).

Tpms function in various F-actin networks that play important roles in cell mechanics, including stress fibers in mammalian cells (Bryce et al., 2003; Percival et al., 2000; Schevzov et al., 2005b; Schevzov et al., 2011; Temm-Grove et al., 1998; Tojkander et al., 2011), F-actin cables in budding and fission yeast (Alioto et al., 2016; Clayton et al., 2014; Liu and Bretscher, 1989) and the contractile ring during cell division in yeast and metazoan cells (Balasubramanian et al., 1992; Eppinga et al., 2006; Hughes et al., 2003; Skau et al., 2009; Stark et al., 2010). To assemble distinct F-actin networks, cells express a complement of Tpm isoforms that can recruit different actin-binding proteins to growing filaments to create specialized networks. In cultured B35 rat cortical neuronal cells, Tpm3.1 recruits myosin II to stress fibers leading to decreased lamellipodia formation and cell migration (Bryce et al., 2003; Schevzov et al., 2005a), while Tpm1.12 (aka TmBr3) increases lamellipodia and cell migration with a reduction in stress fibers (Bryce et al., 2003). In U2OS human osteosarcoma cells, Tpm2.1 (Tm1), Tpm3.1 and Tpm 3.2 stabilize F-actin in focal adhesions while Tpm1.6 and/or Tpm1.7 are needed for dorsal stress fibers (Tojkander et al., 2011). Tpm4.2 recruits myosin II to stress fibers and creates contractile actomyosin bundles by assembling Tpm/myosin II-coated F-actin with branched Arp2/3-nucleated and α-actinin-crosslinked filaments (Tojkander et al., 2011).

While *in vitro* biochemistry and cellular studies point to the importance of specific Tpm isoforms, little is known about which Tpms are needed for assembly and functions of specialized F-actin-networks *in vivo* in non-muscle cells or tissues. Studies of striated muscle demonstrate crucial roles Tpms play in F-actin stabilization in thin filaments, mediating calcium-regulated muscle contraction and proper cycling of myosin cross-bridges (Michele et al., 1999; Ottenheijm et al., 2011). Mutations in Tpms cause human myopathies leading to progressive muscle weakness or hypercontractile muscles (Clarke et al., 2008; Kee and Hardeman, 2008; Marttila et al., 2014; Ochala, 2008; Tajsharghi et al., 2012; Wattanasirichaigoon et al., 2002), indicating that muscle-specific Tpms are critical for force generation and muscle mechanical function. Considerably less is known about the roles of non-muscle Tpms in F-actin networks that contribute to *in vivo* mechanical stability of non-muscle cells or bulk tissue mechanics.

The eye lens is a unique, non-connective tissue where organ function (fine focusing of light entering the eye) requires specific biomechanical properties, presenting an opportunity to study the role of non-muscle Tpms in tissue mechanical functions. The lens is a highly organized, transparent and ellipsoid-shaped organ in the anterior chamber of the eye that fine focuses light onto the retina to transmit a clear image. Bulk lens shape change through tension from ciliary muscles and zonular fibers allows for focusing of near and far objects in a process known as accommodation (Glasser, 2008; Keeney et al., 1995; Millodot, 2009). Age-related lens stiffening causes presybopia, an inability of the lens to change shape to focus on near objects, and the subsequent need for reading glasses. Many studies have documented increases in mouse and human lens stiffness with age (Baradia et al., 2010; Cheng et al., 2016a; Glasser and Campbell, 1998; Glasser and Campbell, 1999; Scarcelli et al., 2011), but the molecular and cellular mechanisms that influence lens biomechanical properties remain unclear.

While our previous work shows that the actin cytoskeleton contributes to lens stiffness, (Cheng et al., 2016b; Gokhin et al., 2012), the Tpm isoform composition of the lens is not known and a role for Tpms in the biomechanical properties of the lens has not been studied. In this study, we perform a comprehensive analysis of all Tpms expressed in mouse lenses and reveal that Tpm3.5 (Tm5NM5) (Geeves et al., 2015) is the major mouse lens Tpm isoform, and is required for normal lens stiffness and resilience. Tpm3.5 is associated with F-actin-rich fiber cell membranes in distinct small puncta that are colocalized with Tmod1. Decreased levels of Tpm3.5 in fiber cells results in dissociation of Tmod1 from the cell membrane and a reorganization of the spectrin-F-actin and of α-actinin-F-actin networks. We conclude that Tpm3.5 promotes Tmod1 association with the spectrin-F-actin network as well as with α-actinin-F-actin networks at fiber cell membranes, and that decreased Tpm3.5 levels with dissociation of Tmod1 lead to a rearrangement of F-actin networks that are then unable to support mechanical load and resilience in lens fiber cells. These data provide the first evidence that any Tpm is required for biomechanical properties in a load-bearing non-muscle tissue and suggests that the Tpm3.5 isoform functions to stabilize mechanical-load-bearing F-actin in non-muscle tissues.

## Results

### Tpm3.5 is the major TM isoform in mouse lenses

To study Tpm function in the lens, we examined *Tpm3/Δexon9d^−/−^* mice, an isoform-specific knockout of exon 9d in *Tpm3* (Fath et al., 2010; Hook et al., 2011; Lees et al., 2013). We first performed RT-PCR and sequencing for Tpm3.1 and Tpm3.2, both containing exon 9d, in the lens. Unexpectedly, we did not detect either Tpm3.1 or Tpm3.2 in the lens (Figure 1B). Brain RNA from *Tpm3/Δexon9d^+/+^* and *Tpm3/Δexon9d^−/−^* mice was used as a control to ensure that the PCR protocol worked and to verify that exon 9d was not expressed in *Tpm3/Δexon9d^−/−^* mice, as expected (Figure 1B). Our previous work had identified a short, Tpm3 exon 9a-containing peptide (aka γTM) in the lens (Nowak et al., 2009). Based on this information, we designed primers to look for transcripts of *Tpm3* containing exon 9a. This analysis revealed that the mouse lens expresses Tpm3.5, a *Tpm3* isoform with exon 9a, and that there appeared to be a slight decrease in Tpm3.5 transcript levels in the lens RNA sample from the *Tpm3/Δexon9d^−/−^* vs. *Tpm3/Δexon9d^+/+^* mice (Figure 1B). Due to high sequence homology between various Tpm isoforms and the large number of splicing variants of each gene, it is difficult to design specific primers for real-time PCR. Thus, in order to verify and quantify the change in transcript level, we performed semi-quantitative PCR for Tpm3.5 and G3DPH (GAPDH, housekeeping gene) with RNA isolated from three pairs of *Tpm3/Δexon9d^+/+^* and three pairs of *Tpm3/Δexon9d^−/−^* lenses. This demonstrated that there was a statically significant decrease in Tpm3.5 transcripts in the *Tpm3/Δexon9d^−/−^* lenses (Figure 1C). We next examined Tpm3.5 protein levels in control and mutant lenses by western blotting using an exon 9a-specific antibody and observed a dramatic decrease in Tpm3.5 levels in the *Tpm3/Δexon9d^−/−^* lenses (Figure 1D). Thus, we conclude that in adult mouse lenses, deletion of exon 9d from the *Tpm3* gene results in an unexpected knockdown of exon 9a-containing Tpm3.5 transcript and protein levels.

**Figure 1.**
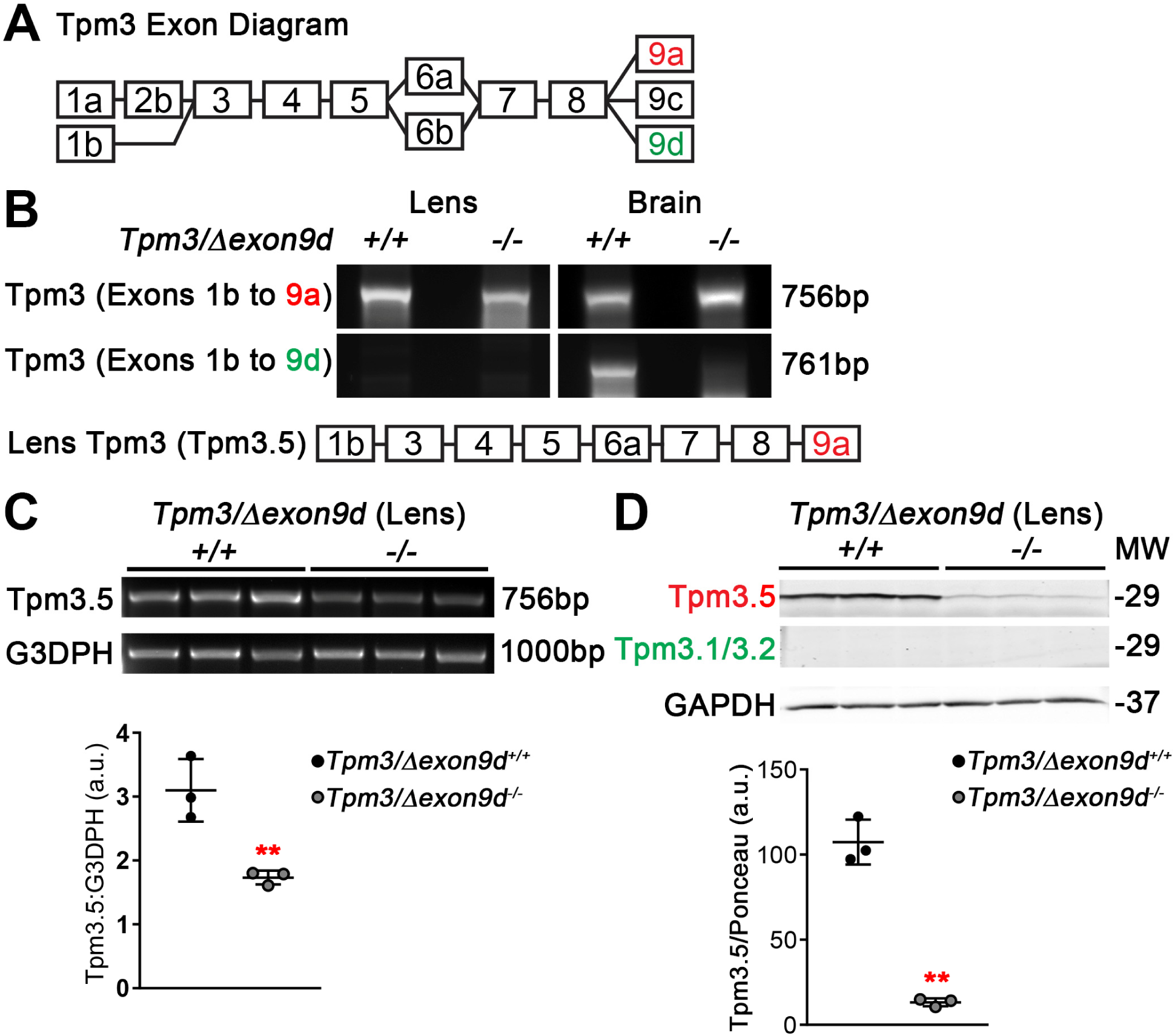
Tpm3.5 is the major lens Tpm isoform. **(A)**Diagram of Tpm3 exons. Alternative splicing produces 6 known Tpm3 isoforms in mice. Not drawn to scale. **(B)**RT-PCR for Tpm3 isoforms in cDNA from *Tpm3/Δexon9d^+/+^* and *Tpm3/Δexon9d^−/−^* lenses and brain reveal that Tpm3-exon 9d is not detected in the lens, and instead contain Tpm3-exon 9a whose levels are unexpectedly reduced in the 9d knockout lens. Brain samples confirm that exon 9d is deleted. **(C)**Semiquantitative RT-PCR from three separate lens *Tpm3/Δexon9d^+/+^* and *Tpm3/Δexon9d^−/−^* cDNA samples reveals decreased *Tpm3.5* transcript levels. G3DPH was used a housekeeping gene and loading control. **(D)**Western blots of *Tpm3/Δexon9d^+/+^* and *Tpm3/Δexon9d^−/−^* lenses reveal significantly decreased levels of Tpm3. 5. Tpm3.1/3.2, containing exon 9d, is not detected in lens samples. GAPDH is shown here to demonstrate equal loading of the samples. Ponceau S staining of total proteins on blots was used as a loading control. Plots reflect mean± SD of n = 3 independent samples per genotype* *, *p<0.01.*

Based on the known isoforms of mouse Tpms (Geeves et al., 2015), we also designed specific primer sets to determine whether other Tpms are expressed in adult mouse lenses, and whether expression of any of these Tpms might be altered in the *Tpm3/Δexon9d^−/−^* mouse lens (Figure S2A). Our RT-PCR and sequencing experiments revealed 5 other Tpm transcripts, Tpms 1.7 (Tm3), 1.8 (Tm5a), 1.9, 1.13 and 4.2 (Table 1 and Figure S2B). These transcripts were present at lower levels than Tpm3.5, and there did not appear to be obvious compensation by altered expression of any of these other Tpm isoforms in *Tpm3/Δexon9d^−/−^* lenses (Figure S2B). We also did not detect any Tpms that are only expressed in *Tpm3/Δexon9d^−/−^* lenses (data not shown). Thus, the *Tpm3/Δexon9d^−/−^* mouse lens provides a tool to understand the function of Tpm3.5.

**Table 1.**
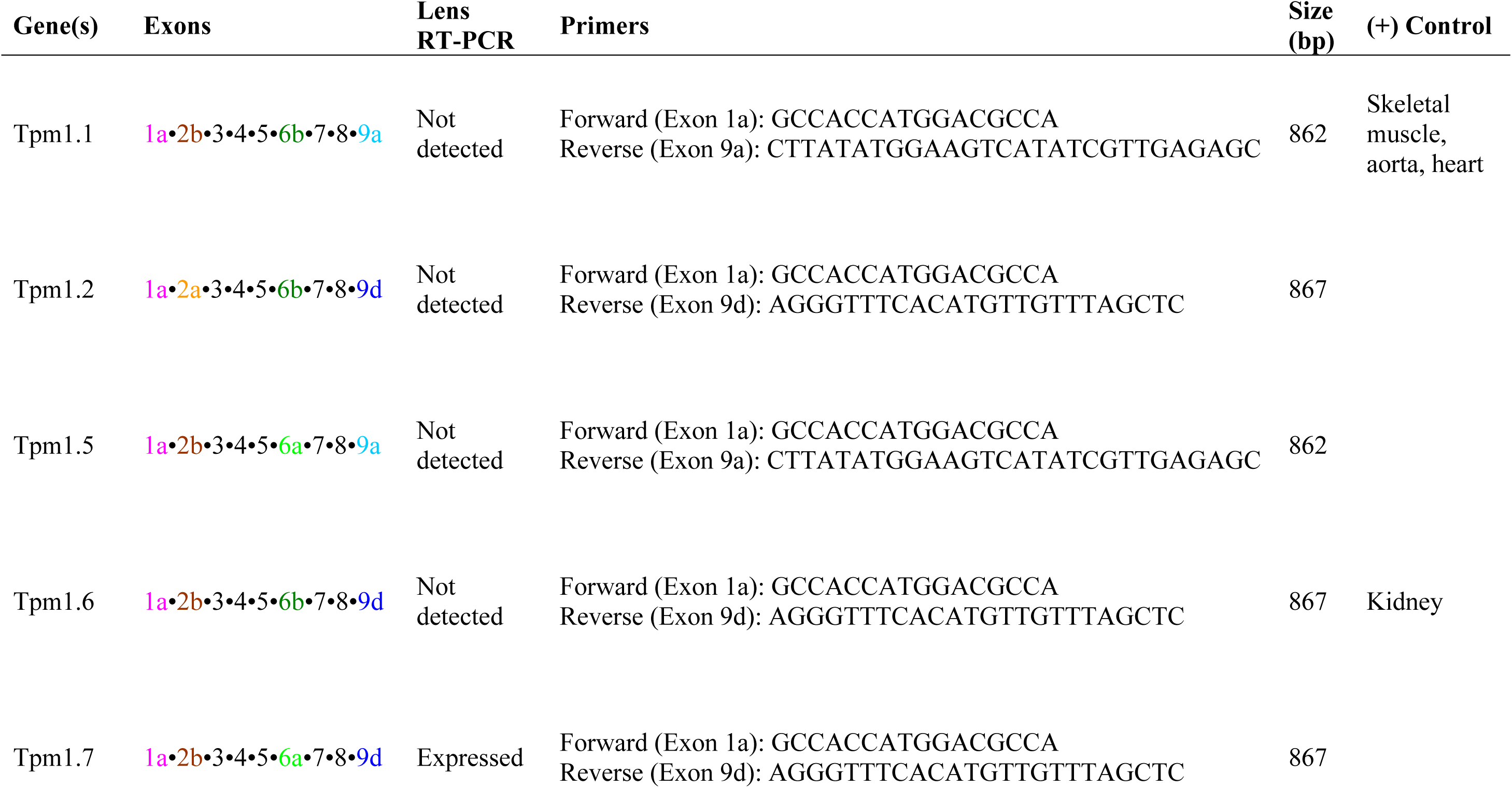
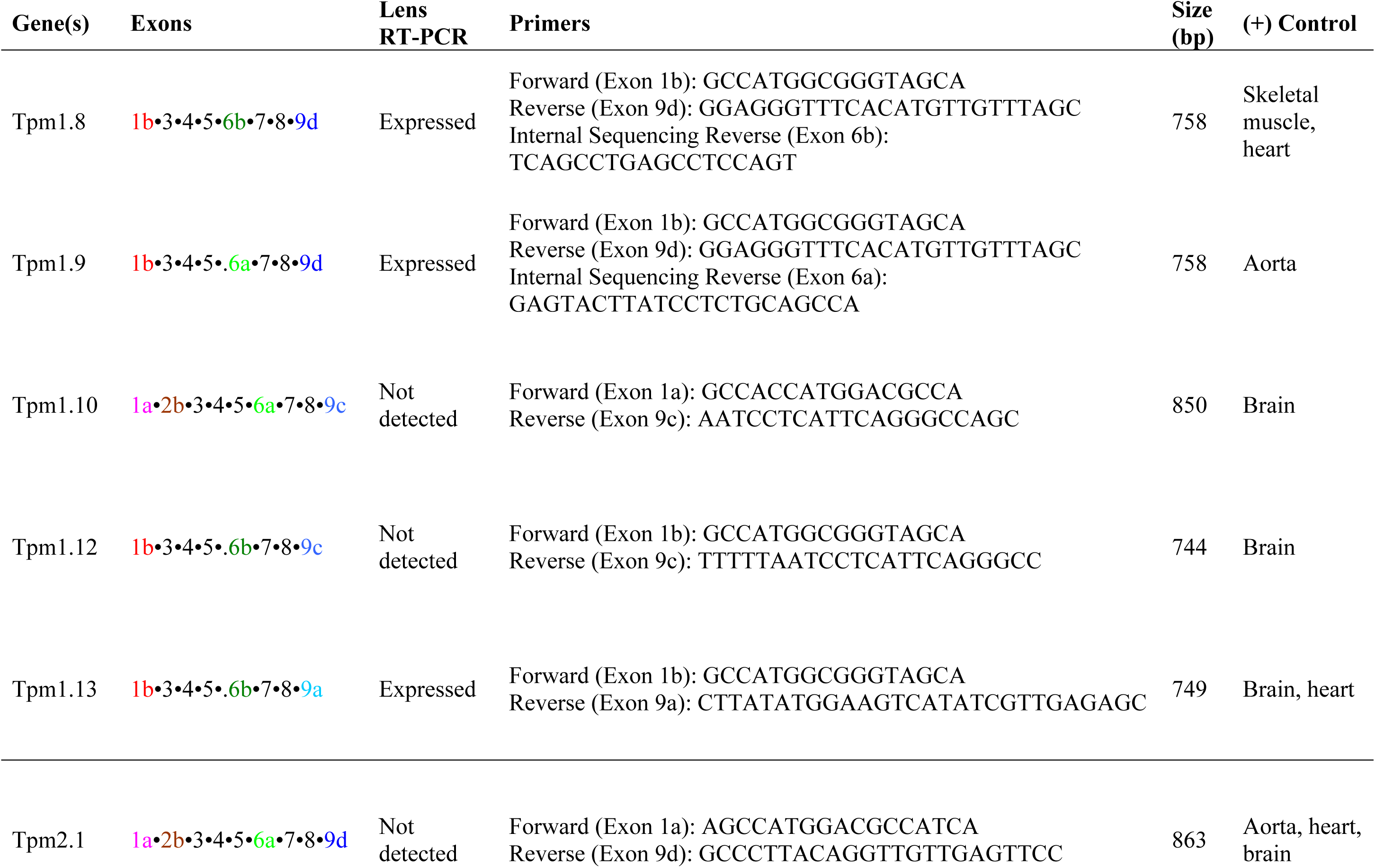
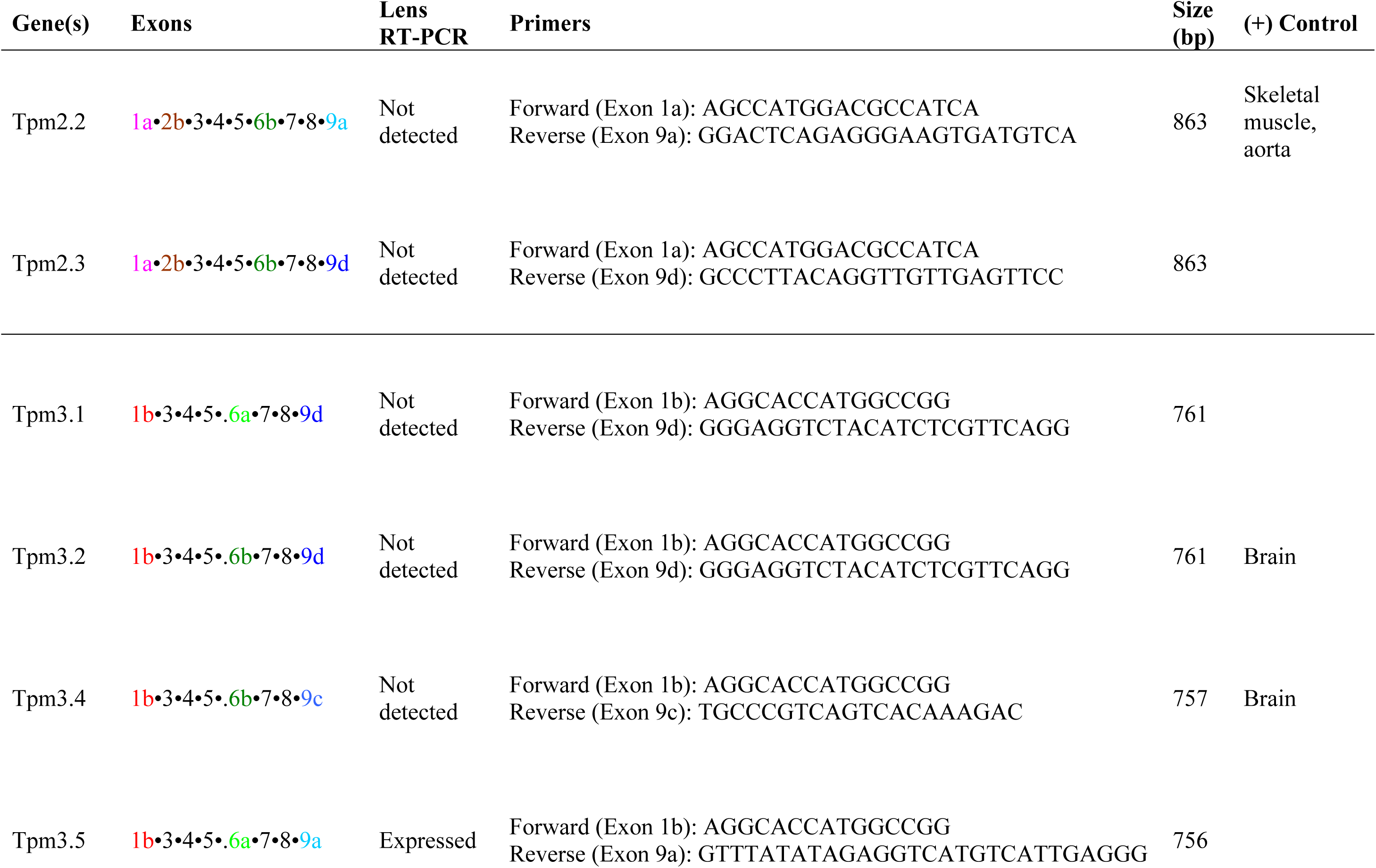
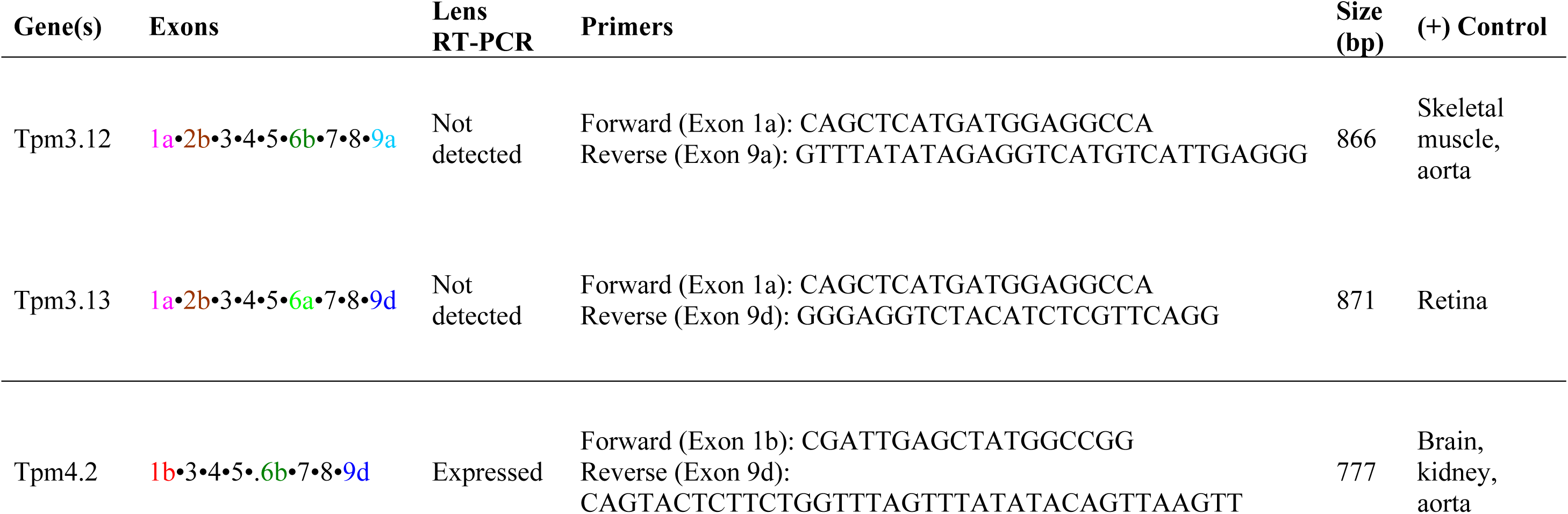
Tpm RT-PCR primers and lens expression data. Primers for each Tpm isoform were designed to cover the entire sequence from exon 1 to exon 9. Since primer pairs may amplify multiple isoforms that differ in exon 2 and/or exon 6, sequencing was used to determine the specific Tpm isoform expressed in lens and other tissues. For Tpms not expressed in the lens, positive (+) control tissue(s) were used to verify the PCR protocol. Four Tpm isoforms were not found in any of the tissues tested, but those primer pairs were verified by a positive control in an alternatively spliced isoform.

### Decreased levels of Tpm3.5 lead to mild anterior cataracts, subtle changes in lens shape, and a marked enlargement of the lens nucleus

To evaluate whether reduced levels of Tpm3.5 lead to alterations in whole lens properties, we first examined 6-week-old lenses from *Tpm3/Δexon9d^+/+^* and *Tpm3/Δexon9d^−/−^* mice and found no obvious cataracts in *Tpm3/Δexon9d^−/−^* lenses (Figure 2A). While we observed sporadic, very small punctate opacities near the anterior pole of *Tpm3/Δexon9d^−/−^* lenses (data not shown), these minor defects occurred at very low frequency and were not further investigated. We also found that while the overall lens volume is unchanged in 8-week-old *Tpm3/Δexon9d^−/−^* lenses, *Tpm3/Δexon9d^−/−^* lenses had a slightly more spherical shape, due to decreased equatorial diameter and increased axial diameter, i.e., decreased aspect ratio (Figure 2C). Subtle anterior opacities and altered lens shape may be due to abnormalities in lens epithelial cell homeostasis or differentiation to fiber cells at the equator, which we have not studied further. Mouse lenses have a very hard central portion where the oldest fiber cells are located, termed the nucleus, that can be mechanically separated from softer cortical fiber cells (Cheng et al., 2016a; Gokhin et al., 2012). Compared to littermate controls, *Tpm3/Δexon9d^−/−^* lenses had increased nuclear volume that occupied a higher fraction of the lens (Figure 2C). These results suggest that Tpm3.5 is important for lens shape and fiber cell maturation.

**Figure 2.**
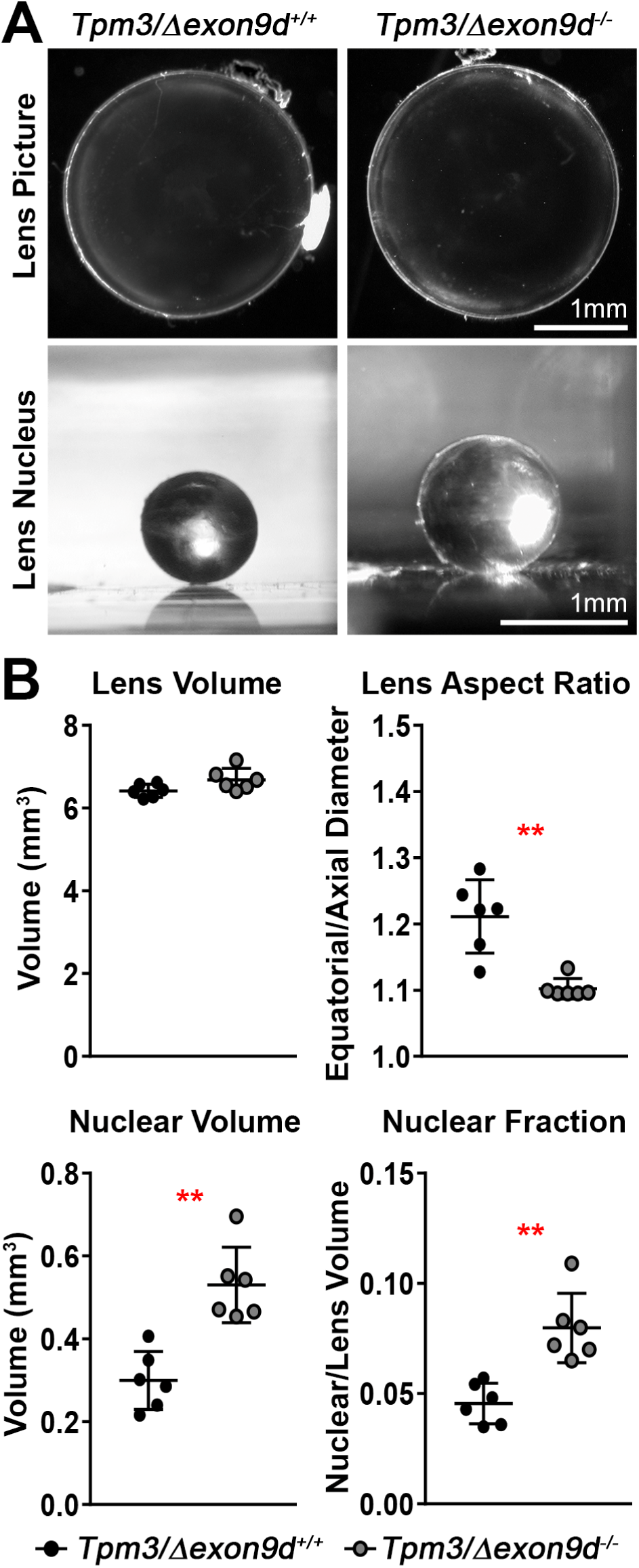
*Tpm3/L1exon9a^−/−^* lenses have mildly altered lens shape and an enlarged lens nucleus. **(A)** Pictures of freshly dissected 6-week-old *Tpm3/Δexon9cf^+/+^* and *Tpm3/Δexon9d^−/−^* lenses. *Tpm3/Δexon9d^−/−^* lenses do not have obvious cataracts. **(B)** The rigid lens nucleus can be dissected away from the soft lens cortex. In the 8-week-old *Tpm3/Δexon9d^−/−^* lens, the nucleus is larger than that in the control lens. **(C)** Morphometric analysis of 8-week-old *Tpm3/Δexon9d^+/+^* and *Tpm3/Δexon9d^−/−^* lenses reveals no change in the lens volume, but *Tpm3/Δexon9d^−/−^* lenses are more spherical with decreased lens aspect ratio. Mutant lenses also had increased nuclear volume and nuclear fraction. Plots reflect mean ± SD of n = 6 lenses per genotype. **, *p*<0.01. Scale bars, 1mm.

### Tpm3.5 is required for lens stiffness and resilience

To determine whether decreased Tpm3.5 leads to altered lens biomechanical properties, we measured the stiffness of 8-week-old *Tpm3/Δexon9d^+/+^* and *Tpm3/Δexon9d^−/−^* lenses using a simple coverslip compression method (Cheng et al., 2016a; Gokhin et al., 2012). In this method, the lens is viewed through a dissection microscope and coverslips are sequentially loaded onto the tissue resulting in axial compression and equatorial expansion of the lens. Images taken of the lens under load were then used to calculate percent change in axial and equatorial dimensions (strain) and recovery after removal of load (resilience). In *Tpm3/Δexon9d^−/−^* lenses, there was an increase in axial compressive strain at high loads (7-10 coverslips) as well as in equatorial strain at high loads (9-10 coverslips) (Figure 3A and 3B), indicating that mutant lenses are softer than control lenses at high mechanical loads. Dot plots of axial and equatorial strains show no detectable difference in the stiffness of mutant lenses at low load (1 coverslip) while there is a significant increase in strain at the maximum load (10 coverslips), indicative of softer lenses (Figure 3C and 3D). We also examined recovery of lens shape after removal of load by calculating the ratio between the pre-loading and post-loading axial diameter (resilience). Our data revealed that *Tpm3/Δexon9d^−/−^* lenses had dramatically decreased resilience and only recovered to 88.86% ± 2.03% of the pre-loading axial diameter (Figure 3E and 3F). These data indicate that Tpm3.5 is needed for normal lens stiffness and resilience. We also conclude that the hard lens nucleus does not appear to affect bulk lens stiffness in an axial compression assay, since decreased Tpm3.5 leads to a softer lens despite the larger size of the hard central lens nucleus.

**Figure 3.**
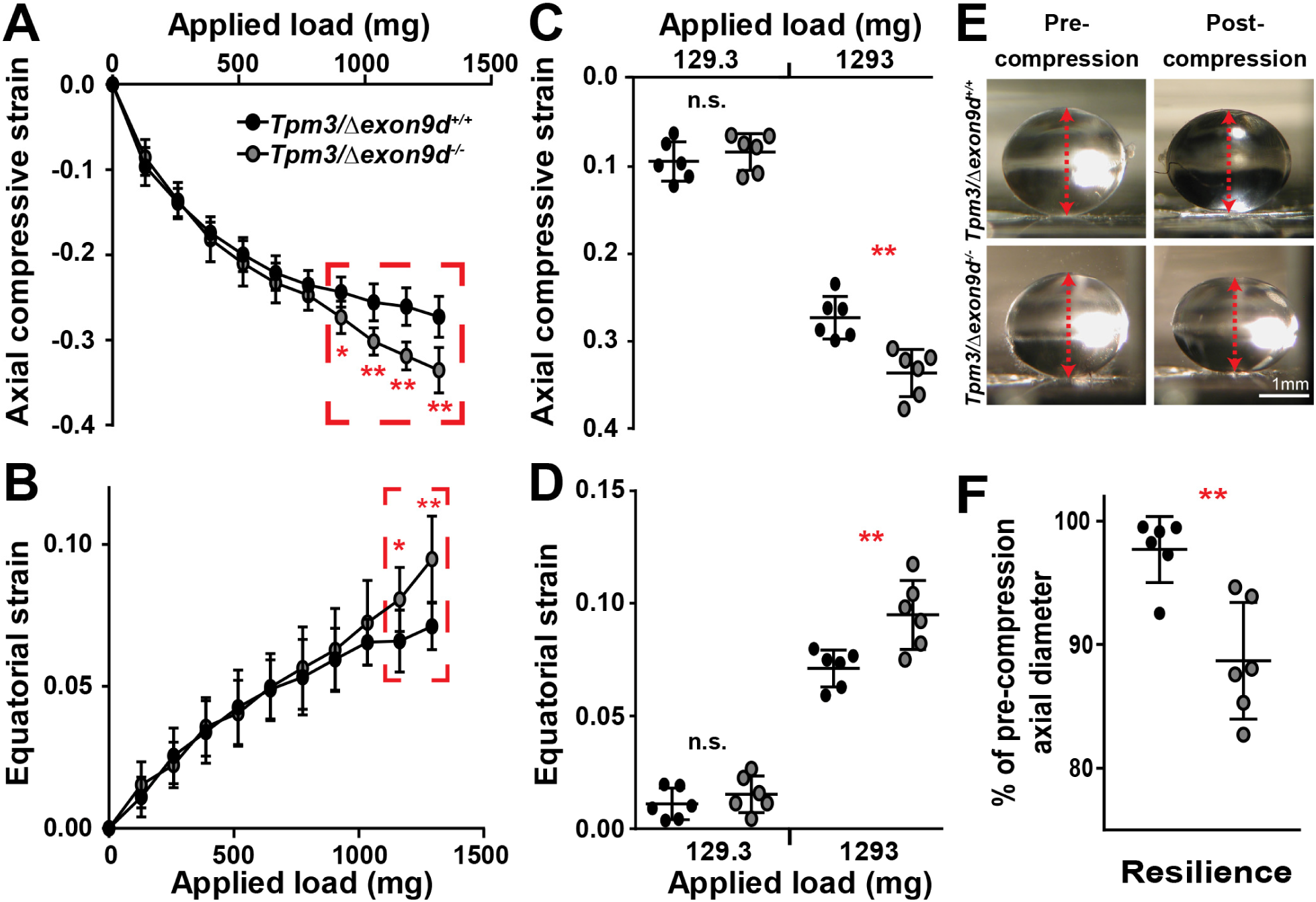
*Tpm3/Δexon9d ^−/−^* lenses are softer at high mechanical loads and have decreased resilience after compression. **(A-B)** Compression testing of 8-week-old *Tpm3/Δexon9d^+/+^* and *Tpm3/Δexon9d^−/−^* lenses revealed increased axial and equatorial strain at high mechanical loads (1034.4mg-1293mg, 8-10 coverslips). **(C-D)** Dot plots of axial and equatorial strain at the lowest load (129.3mg, 1 coverslip) and the maximum load (1293mg, 10 coverslips) showed increased strain in *Tpm3/Δexon9d^−/−^* lenses at the maximum load. **(E)** Side view pictures of *Tpm3/Δexon9d^+/+^* and *Tpm3/Δexon9d^−/−^* lenses pre-compression and post-compression. The red dashed lines indicate the axial diameter. **(F)** *Tpm3/Δexon9d^−/−^* lenses had decreased resilience, calculated as the ratio of the pre-compression over post-compression axial diameter. While control *Tpm3/Δexon9d^+/+^* lenses recovered to 97.24% ± 0.83% of the pre-compression axial diameter, mutant *Tpm3/Δexon9d^−/−^* lenses only returned to 88.86% ± 2.03% of the pre-loading axial diameter. This indicates that mutant lenses were unable to recover their normal shape after removal of mechanical load. Plots reflect mean± SD of n = 6 lenses per genotype. *, *p*<0.05; **, *p*<0.01. Scale bar, lmm.

### Tpm3.5 is associated with F-actin-rich fiber cell membranes, but decreased levels of Tpm3.5 do not affect fiber cell hexagonal packing nor intercellular interdigitations

Cells in the lens are encapsulated by a thin collagenous basement membrane called the lens capsule. A monolayer of epithelial cells covers the anterior hemisphere, and the bulk of the lens is composed of elongated fiber cells (Figure S1A). Fibers are long and thin cells that stretch from the anterior to posterior poles. Fiber cells are hexagonal in cross section to effectively pack cells and minimize light scattering intercellular spaces (Kuszak et al., 2006; Kuszak et al., 2004). We examined the packing of hexagonal fiber cells in *Tpm3/Δexon9d^+/+^* and *Tpm3/Δexon9d^−/−^* lenses by immunostaining lens frozen sections in the cross orientation. Staining for Tpm3.5 and F-actin reveals organized cortical fiber cells packed into neat rows of hexagonal cells in both *Tpm3/Δexon9d^+/+^* and *Tpm3/Δexon9d^−/−^* lenses, indicating that decreased Tpm3.5 does not affect overall fiber cell shape and hexagonal packing organization (Figure 4). Tpm3.5 colocalizes with F-actin along fiber cell membranes of *Tpm3/Δexon9d^+/+^* lenses, in agreement with previous work (Nowak et al., 2009), and is also present in the lens epithelium. Lower Tpm3.5 staining signal is observed in mutant lens sections, as expected from the western blots (Figure 1D). Further, while F-actin staining appears to be crisply delineated at the mutant lens fiber cell membrane, Tpm3.5 staining appears more cytoplasmic (Figure 4, asterisks).

**Figure 4.**
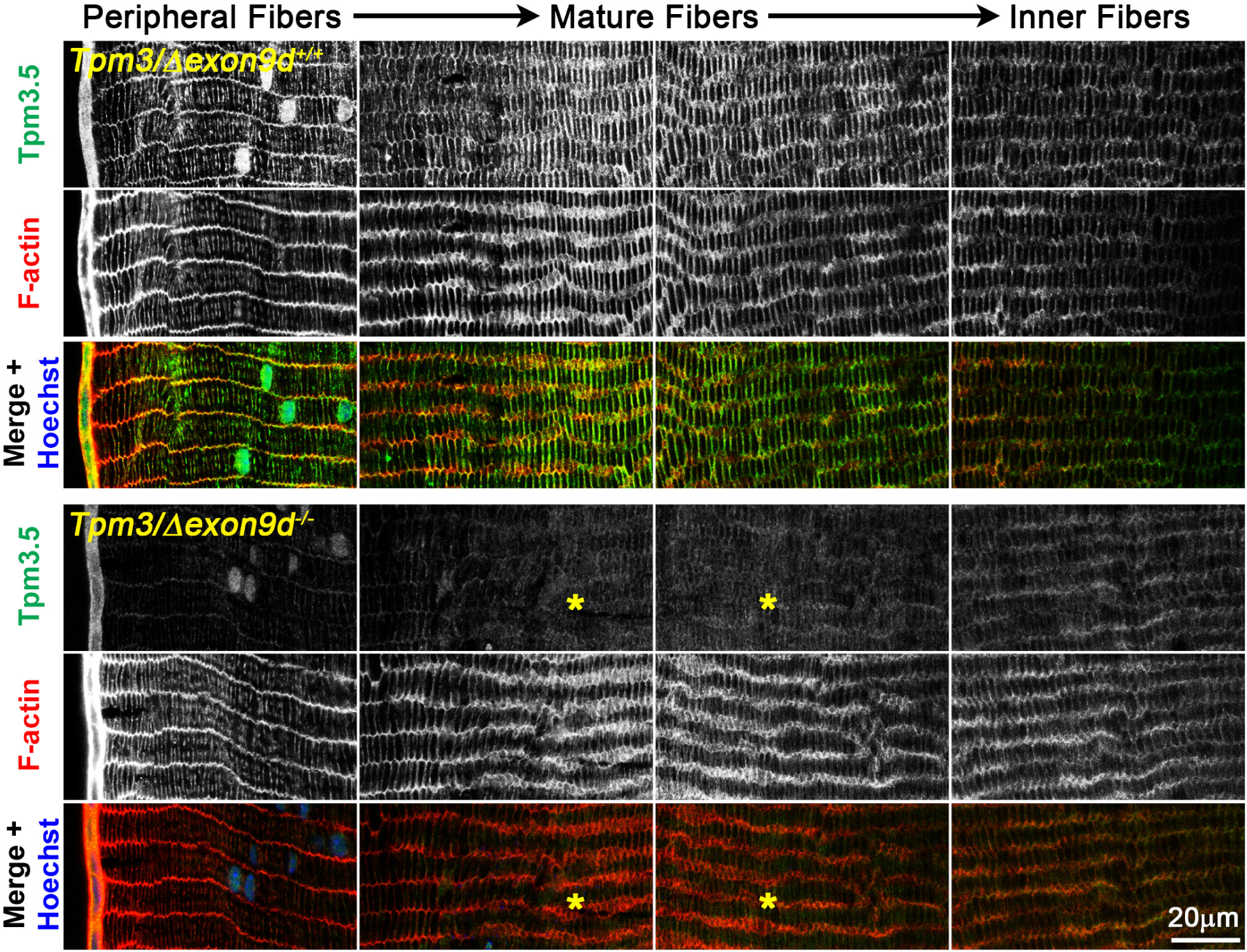
Decreased level of Tpm3.5 does not cause obvious changes in lens fiber cell packing. Immunostaining of frozen sections in the cross orientation from *Tpm3/Δexon9d^+/+^* and *Tpm3/Δexon9d^−/−^* lenses for Tpm3.5 (green), F-actin (red) and cell nuclei (Hoechst, blue). Images are from sections near the lens equator with panels from the lens periphery (left most) toward the inner fibers (right most). There is no obvious change in hexagonal fiber cell packing in the *Tpm3/Δexon9d^−/−^* lens section. However, there is a decrease in Tpm3.5 staining signal and areas where the Tpm3.5 staining signal appears abnormally cytoplasmic in the mutant lens section (asterisks). Scale bar, 20µm.

During cell maturation, lens fibers undergo a unique process to remove all cellular organelles and develop specialized F-actin-rich interlocking membrane interdigitations. Peripheral cortical lens fiber cells are relatively straight with small F-actin rich protrusions (~1µm long) along their lengths while mature fibers have an undulating morphology with small F-actin-rich protrusions projecting from large F-actin-containing paddle domains (~5-10µm) (Figure S1) (Blankenship et al., 2007; Cheng et al., 2016b; Kuwabara, 1975; Lo et al., 2014; Willekens and Vrensen, 1981; Willekens and Vrensen, 1982). It has long been hypothesized that these interlocking domains are required for mechanical integrity during lens shape change. Therefore, we examined Tpm3.5 and F-actin subcellular localization with respect to fiber cell membrane morphologies in individual lens fiber cells from *Tpm3/Δexon9d^+/+^* and *Tpm3/Δexon9d^−/−^* lenses, isolated as previously described (Cheng et al., 2016b). Due to the small sizes of the F-actin-rich membrane protrusions and the complex geometry of mature fiber cells, we performed super-resolution confocal microscopy using Airyscan, with a resolution of 140nm laterally and 400nm axially. Single optical sections through the center of the cell are presented in all individual fiber cell immunostaining images. Tpm3.5 is localized in small puncta in F-actin-rich regions along the membrane of mature lens fiber cells (Figure 5), and Tpm3.5 staining signal is dramatically reduced in mutant lens fibers (Figure 5), as expected (Figure 1D and Figure 4). Enlargements of control *Tpm3/Δexon9d^+/+^* mature fiber cell images reveal that Tpm3.5 puncta are associated with F-actin in the small protrusions and at their bases (Figure 5B, arrows). In contrast, in the mutant *Tpm3/Δexon9d^−/−^* mature lens fiber, small Tpm3.5 puncta no longer appear in protrusions, and only a few residual Tpm3.5 puncta are observed near the base of small protrusions (Figure 5B, open arrowheads). Despite reduced overall levels of Tpm3.5 and its selective absence from small protrusions, no obvious changes in F-actin distribution or intensity are apparent in mutant *Tpm3/Δexon9d^−/−^* lens fibers. Moreover, unlike the loss of large paddles in mature *Tmod1^−/−^* lens fibers (Cheng et al., 2016b), there are no detectable differences in the morphologies of large paddles or small protrusions in the *Tpm3/Δexon9d^−/−^* lens fibers.

**Figure 5.**
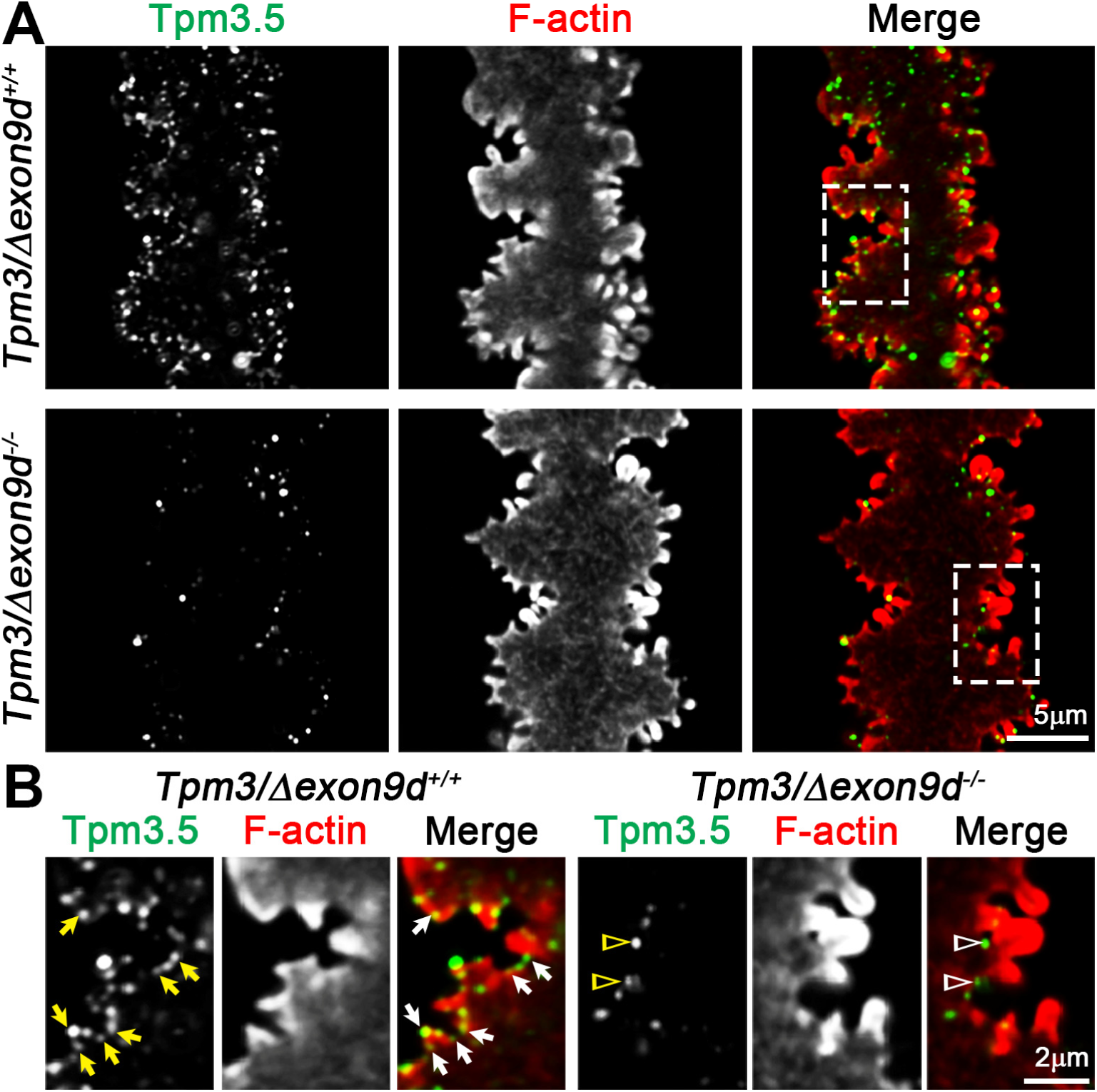
Tpm3.5 is localized in small puncta along the fiber cell membrane. Immunostaining of single fiber cells from 6-week-old *Tpm3/Δexon9d^+/+^* and *Tpm3/Δexon9d^−/−^* lenses for Tpm3.5 (green) and F-actin (red). Images are single optical sections through the cell cytoplasm. **(A)** In the mature *Tpm3/Δexon9d^+/+^* fiber cell, Tpm3.5 is enriched in puncta along the cell membrane, colocalizing with F-actin. There is an obvious decrease in Tpm3.5 staining in the mature *Tpm3/Δexon9d^−/−^* lens fiber. The mutant mature fiber has large paddle domains decorated by small protrusions similar to the control lens fiber. **(B)** An enlargement of a paddle region from the control mature fiber reveals Tpm3.5 puncta along the cell membrane in at the tips of and near the base of small protrusions (arrows). In the mutant lens fiber, Tpm3.5 staining is drastically reduced at the cell membrane and remaining Tpm3. 5 small puncta are mostly at the base of small protrusions (open arrowheads). Scale bars, 5µm in A and 2µm in B.

To verify that there is no change in complex interdigitations between mutant lens fibers, we also performed SEM of microdissected lens halves to visualize the fiber cell profiles *in situ* in 2-month-old *Tpm3/Δexon9d^+/+^* and *Tpm3/Δexon9d^−/−^* lenses. Consistent with single fiber cell immunostaining experiments, there are no obvious differences in small protrusions or large paddle domains between *Tpm3/Δexon9d^+/+^* and *Tpm3/Δexon9d^−/−^* mature lens fibers (Figure 6). We also compared inner fiber cells of the lens nucleus and did not see any obvious changes in those cells. Thus, reduced levels of Tpm 3.5 protein in *Tpm3/Δexon9d^−/−^* lenses do not appear to affect fiber cell morphology, including formation of interlocking interdigitations between lens fibers. Therefore, we conclude that loss of stiffness at high mechanical load and loss of resilience in *Tpm3/Δexon9d^−/−^* lenses are not due to morphological alterations in interlocking interdigitations between fiber cells.

**Figure 6.**
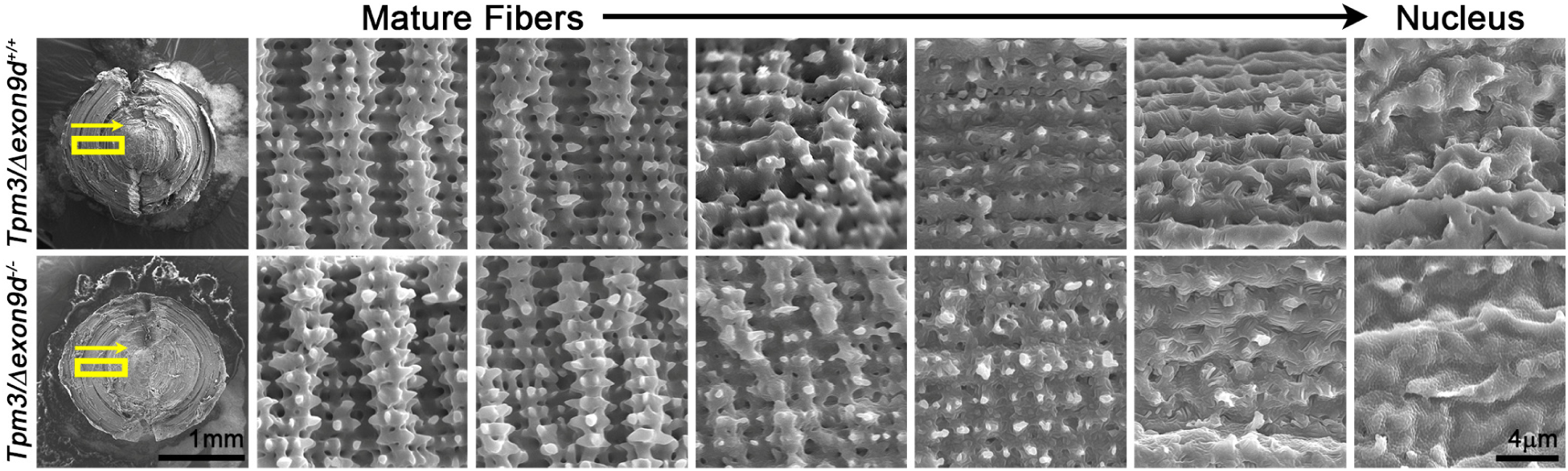
SEM confirms that mutant *Tpm3/Δexon9d^−/−^* lens fibers have normal cell morphology. SEM pictures from 8-week-old *Tpm3/Δexon9d^+/+^* and *Tpm3/Δexon9d^−/−^* lenses taken from maturing lens fibers (left) to lens nucleus (right). Both control and mutant lens fibers have normal paddle domains and small protrusions, and cell membranes remodel to form smoother tongue-and-groove contours in the innermost fiber cells of the lens nucleus. There are no obvious morphological differences between control and mutant lens fibers. Scale bars, 1mm and 4μm.

### Tpm3.5 stabilizes Tmod1 and the β2-spectrin-based membrane skeleton in lens fiber cells

Tpms enhance the capping affinity of Tmods for F-actin pointed ends, and together, Tpms and Tmods promote F-actin stability *in vitro* (Kostyukova, 2008; Lewis et al., 2014; Rao et al., 2014; Yamashiro et al., 2012; Yamashiro et al., 2014), in the spectrin-F-actin membrane skeleton of lens fiber cells (Cheng et al., 2016b; Gokhin et al., 2012) and polarized epithelial cells (Nowak et al., 2009; Weber et al., 2007), and in contractile myofibrils of cardiac myocytes (Cheng et al., 2016b; Gokhin et al., 2012; Mudry et al., 2003). Therefore, we hypothesized that decreased levels of Tpm3.5 in the lens might reduce association of Tmod1 with F-actin, and lead to F-actin and membrane skeleton instability, thereby compromising lens stiffness.

To test this idea, we first determined whether Tpm3.5 is colocalized with Tmod1 and F-actin at the membrane of lens fibers. Double immunolabeling revealed that Tpm3.5 and Tmod1 are often colocalized in small puncta along the membrane of the small F-actin rich protrusions in the control *Tpm3/Δexon9d^+/+^* mature lens fiber cells (Figure 7, open arrowheads in 7B). We had not previously detected Tmod1 along the base of small protrusions (Cheng et al., 2016b), which has now been revealed by highly sensitive super-resolution Airyscan confocal microscopy. In control lens fiber cells, we observe that Tmod1 is enriched in valleys between large paddles (Figure 7, arrows), consistent with our previous data (Cheng et al., 2016b), while Tpm3.5 does not appear to be enriched in the valleys between large paddles. By contrast, in the Tpm3.5-deficient mutant lens fibers, Tmod1 now appears dispersed in many small puncta throughout the cytoplasm (Figure 7A, asterisk and 7B, open arrows), and there is little or no colocalization of residual Tpm3.5 with the remaining small Tmod1 puncta at the membrane (Figure 7B, open arrowheads). Interestingly, residual Tpm3.5 staining is enriched and colocalized with the few remaining large Tmod1 puncta that persist in the valleys between large paddles in the *Tpm3/Δexon9d^−/−^* mature lens fibers (Figure 7B, arrows). Immunostaining of lens cross sections also revealed dissociation of Tmod1 from mature fiber cell membranes in the *Tpm3/Δexon9d^−/−^* sample (Figure S2). We conclude that reduced levels of Tpm3.5 lead to dissociation of Tmod1 from F-actin at the fiber cell membrane. These cytoplasmic Tmod1 puncta may reflect abnormal Tmod1 capping of transcellular F-actin networks that are weakly stained in the fiber cell cytoplasm (Figure 7A, asterisk), as immunostainning of cytosolic Tmod1 would be expected to be diffuse, not punctate.

**Figure 7.**
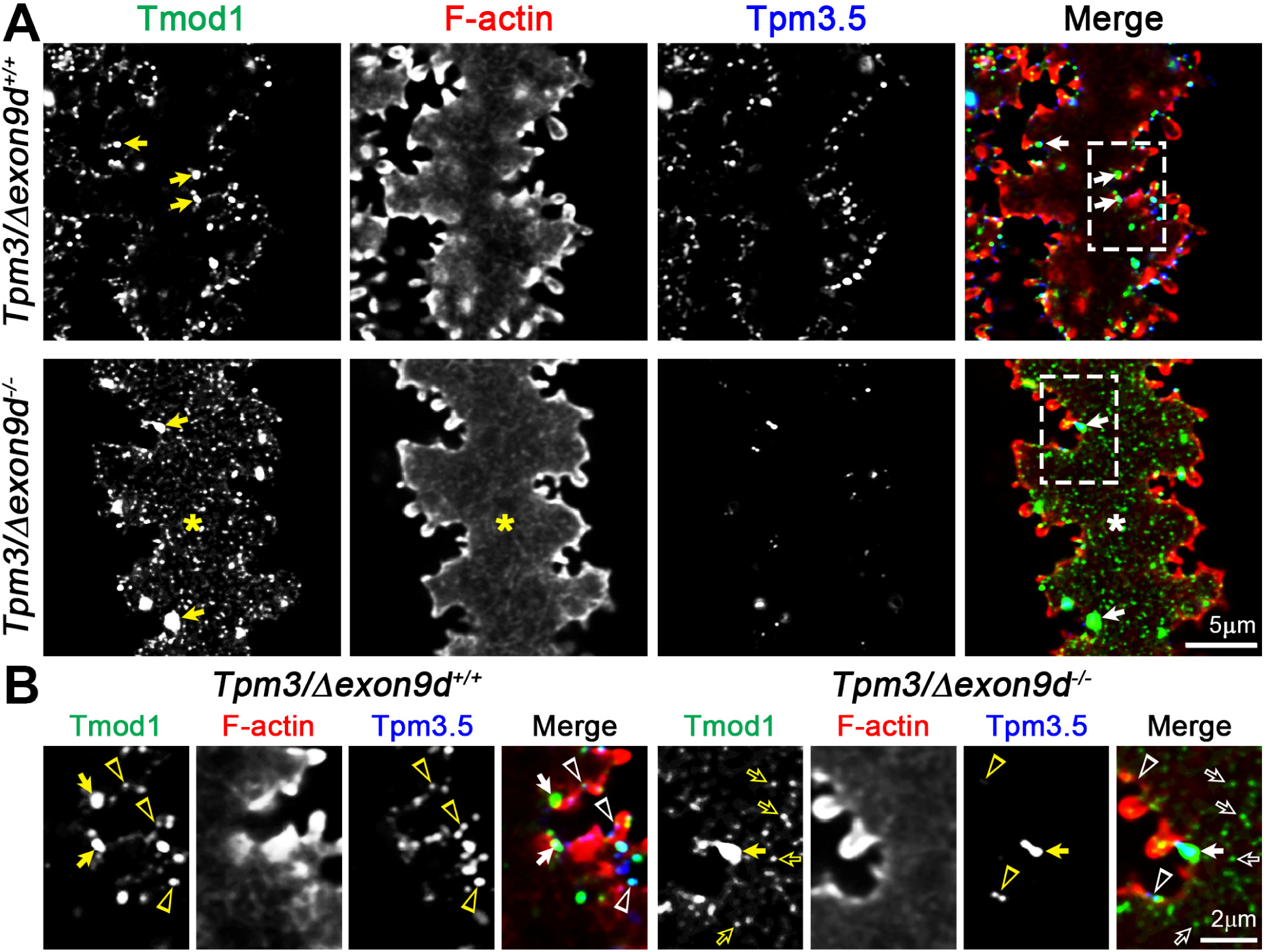
Tmod1 and Tpm3.5 are colocalized with F-actin at the membrane of lens fiber cells. Immunostaining of single mature fiber cells from 6-week-old *Tpm3/Δexon9d^+/+^* and *Tpm3/Δexon9d^−/−^* lenses for Tmod1 (green), F-actin (red) and Tpm3.5 (blue). Images are single optical sections through the cell cytoplasm. **(A)** In the *Tpm3/Δexon9d^+/+^* mature lens fiber, we observe bright Tmod1 puncta in the valleys between large paddles (arrows) and dimmer Tmod1 puncta along with Tpm3.5 puncta along the cell membrane. The mutant mature fiber cell has decreased Tpm3.5 staining along with appearance of Tmod1 puncta in the cell cytoplasm (asterisk). Bright Tmod1 puncta remain at the membrane in the valleys between large paddles in the mutant fiber cell. In the cytoplasm of both control and mutant lens fibers, there is a weakly stained transcellular F-actin network. **(B)** An enlargement from the control lens fiber reveals that Tmod1 and Tpm3.5 puncta are often colocalized along the cell membrane (open arrowheads). While Tmod1 is enriched in the valleys between large paddles (arrows), there is no obvious increase in Tpm3.5 staining in these areas. In the mutant lens fiber, most of the Tpm3.5 signal is absent from the cell membrane with only a few remaining puncta of staining (open arrowheads). Tmod1 puncta are now in the cytoplasm (open arrows), and the residual Tpm3.5 is enriched in the valleys between large paddles along with Tmod1 (arrow). Scale bars, 5µm in A and 2μm in B.

Next, to test the possibility that reduction of Tpm3.5 levels and dissociation of Tmod1 in the *Tpm3/Δexon9d^−/−^* lens might affect membrane skeleton organization, we examined the localization of membrane skeleton component, β2-spectrin, in single lens fibers from *Tpm3/Δexon9d^+/+^* and *Tpm3/Δexon9d^−/−^* lenses. In control lens fiber cells, β2-spectrin colocalizes with Tmod1 in puncta along the F-actin-rich membranes, with especially large bright Tmod1 and β2-spectrin puncta located in the valleys between large paddles (Figure 8, arrows), as previously shown (Cheng et al., 2016b), as well as in small puncta near the base of small protrusions (Figure 8B, arrowheads). By contrast, in mature fibers of *Tpm3/Δexon9d^−/−^* lenses, β2-spectrin staining appears increased and more continuous along the membrane at the bases of small protrusions (Figure 8, open arrowheads), unlike the discrete β2-spectrin puncta in *Tpm3/Δexon9d^+/+^* lens fibers. Again, we observe that Tmod1 staining is in the mutant cell cytoplasm (Figure 8B, open arrows). Thus, decreased levels of Tpm3.5 lead to redistribution of β2-spectrin at the membrane to a more continuous pattern. Other work has shown that Tpms can inhibit the binding of spectrin to F-actin *in vitro* (Fowler and Bennett, 1984b). Thus, we hypothesize that β2-spectrin may now occupy regions of F-actin at the fiber cell membrane that would have been occupied by Tpm3.5, suggesting a change in the organization of the F-actin network.

**Figure 8.**
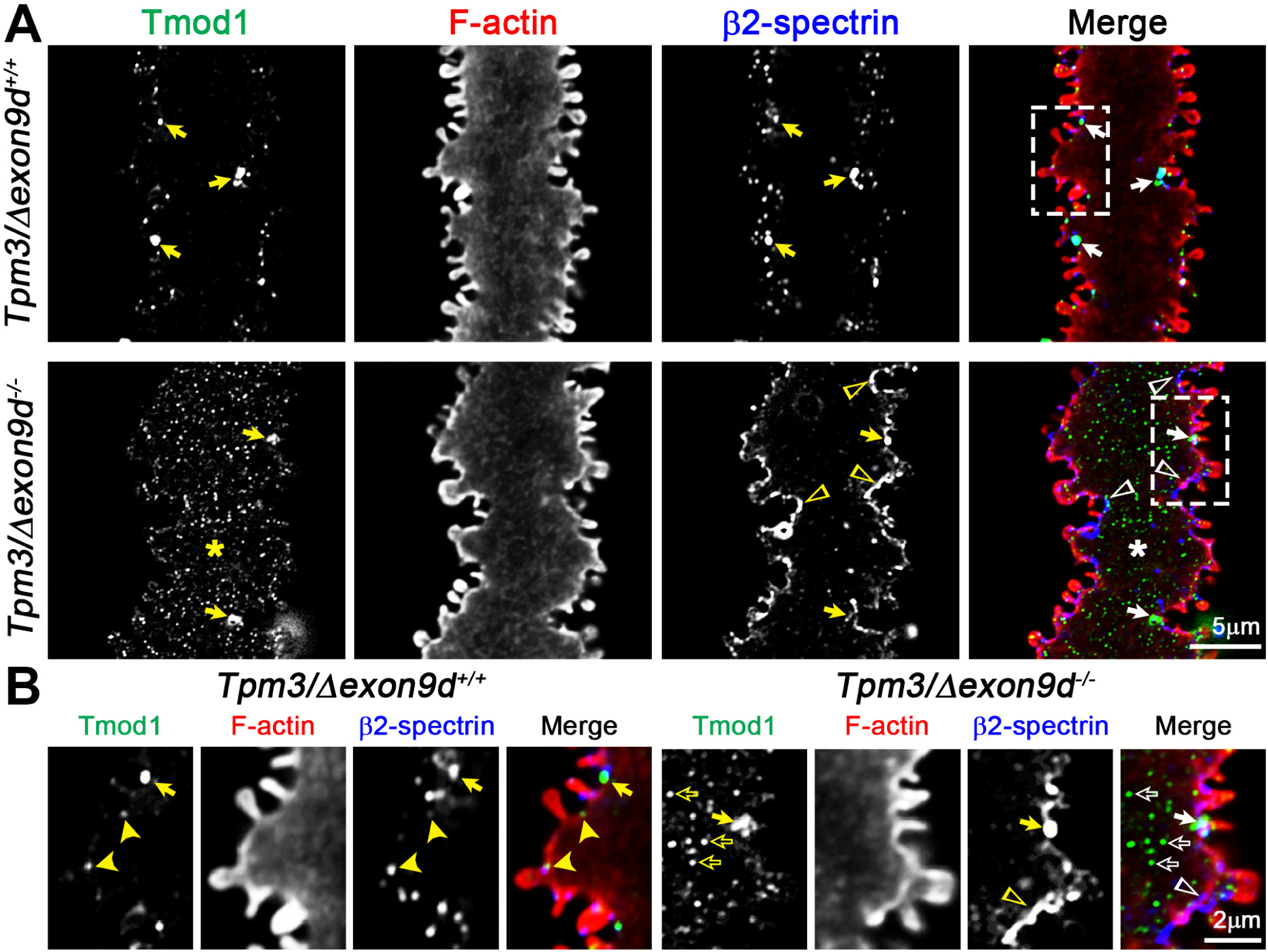
β2-spectrin localization is altered in *Tpm3/Δexon9d^−/−^* lens fiber cells. Immunostaining of single fiber cells from 6-week-old *Tpm3/Δexon9d^+/+^* and *Tpm3/Δexon9d^−/−^* lenses for Tmod1 (green), F-actin (red) and β2-spectrin (blue). Images are single optical sections through the cell cytoplasm. **(A)** In the control mature fiber cell, there are large puncta of Tmod1 and β2-spectrin staining in the valleys between large paddles (arrows). In the mutant mature fiber cells, while large puncta of Tmod1 and β2-spectrin remain in the valleys between large paddles (arrows), most of the Tmod1 staining is now in the cell cytoplasm (asterisk) and β2-spectrin staining is expanded along the cell membrane (open arrowheads). **(B)** An enlargement from the control mature fiber cell reveals Tmod1 puncta in the valley between large paddles (arrow) and along the cell membrane (arrowheads). In the mutant lens fiber, Tmod1 puncta are present in the cytoplasm (open arrows), and there is expanded β2-spectrin staining (open arrowheads). Enriched Tmodl and 2-spectrin can still be found in the valley between large paddles in the mutant fiber cell (arrow). Scale bars, 5µm in A and 2µm in B.

### Localization of α-actinin and fimbrin, but not Arp3 or N-cadherin, is altered in mutant lens fiber cells

To further probe the organization of F-actin networks in Tpm3.5-deficient lens fiber cells, we examined the localization of several F-actin binding proteins, α-actinin, fimbrin and Arp2/3, whose binding to F-actin are all modulated by Tpms (Blanchoin et al., 2001; Brayford et al., 2016; Bugyi et al., 2010; Christensen et al., 2017; Gateva et al., 2017; Hsiao et al., 2015; Skau and Kovar, 2010; Winkelman et al., 2016). In the *Tpm3/Δexon9d^+/+^* mature lens fiber cell, α-actinin puncta are enriched in the valleys between large paddles (Figure 9A, arrows), while α-actinin staining appears increased and dispersed more broadly along the cell membrane in the mutant *Tpm3/Δexon9d^−/−^* fiber (Figure 9A, open arrowheads), similar to β2-spectrin (Figure 8). Since Tpms inhibit α-actinin binding to F-actin, increased α-actinin in Tpm3.5-deficient lenses may be a consequence of newly available binding sites for α-actinin on F-actin at the fiber cell membrane, similar to the mechanism proposed above for increased β2-spectrin.

**Figure 9.**
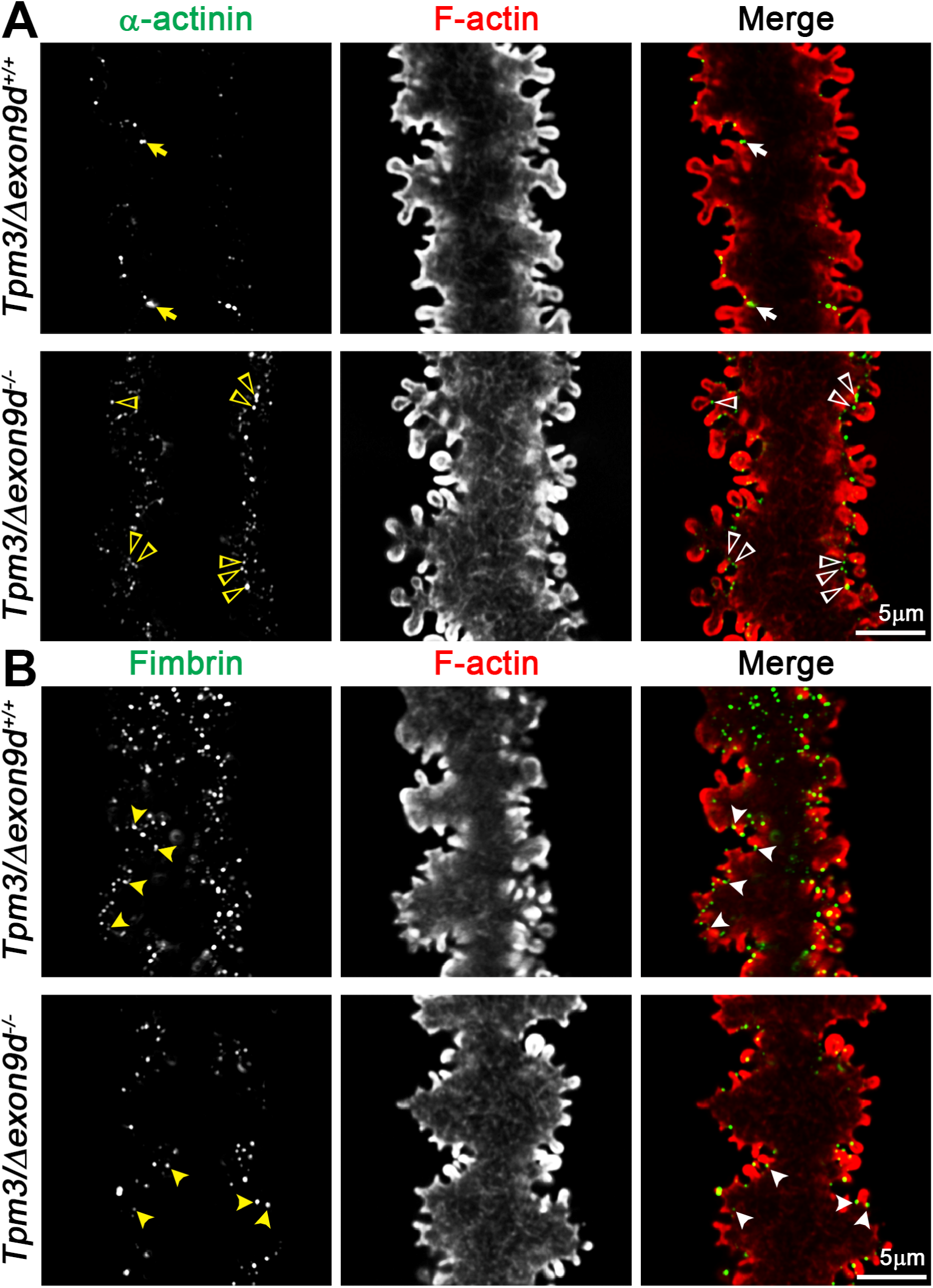
The α-actinin-F-actin network is expanded while fimbrin-bundled F-actin is reduced in *Tpm3/Δexon9d^−/−^* lens fibers. **(A)** Immunostaining of single mature fiber cells from 6-week-old *Tpm3/Δexon9d^+/+^* and *Tpm3/Δexon9d^−/−^* lenses for α-actinin (Actn1, green) and F-actin (red). Images are single optical sections through the cell cytoplasm. In the control lens fiber, α-actinin is enriched in bright puncta in the valleys between large paddles (arrows). In the mutant lens fiber, α-actinin staining is expanded to numerous small puncta along the cell membrane in the valleys between large paddles and at the base of small protrusions (open arrowheads). **(B)** Immunostaining of single mature fiber cells from 6-week-old *Tpm3/Δexon9d^+/+^* and *Tpm3/Δexon9d^−/−^* lenses for fimbrin (plastin, green) and F-actin (red). Fimbrin is localized in small puncta at the base of small protrusions in the control lens fiber (arrows). While the fimbrin localization pattern appears unaffected in the mutant lens fiber (arrows), the numbers of fimbrin puncta associated with the membrane protrusions appear reduced. Scale bars, 5µm.

We also investigated the localization of fimbrin in control and mutant lens fibers. In the control *Tpm3/Δexon9d^+/+^* lens fiber cell, fimbrin is localized to abundant small puncta near the base of small protrusions (Figure 9B, arrows), consistent with our previous data (Cheng et al., 2016b). In contrast to α-actinin staining, which increases in mutant *Tpm3/Δexon9d^−/−^* lens fibers, fimbrin staining is noticeably reduced at the cell membrane, though it still appears punctate (Figure 9B, filled arrowheads). Reduced fimbrin at the mutant cell membrane may be a consequence of the widely spaced filaments characteristic of α-actinin-crosslinked F-actin networks that precludes fimbrin cross-linking of F-actin into compact bundles (Alberts; Vignjevic et al., 2006; Winkelman et al., 2016).

While F-actin cross-linking proteins are affected by decreased Tpm3.5 levels (Figures 8 and 9), we observe no obvious changes in Arp3 or N-cadherin localization between control and mutant mature lens fiber cells (Figure S4). Localization of Arp3 and N-cadherin in small puncta near the base of small protrusions is similar to our previous data (Cheng et al., 2016b). These results suggest that adherens junctions and Arp2/3-nucleated branched F-actin networks are unlikely to be affected by decreased Tpm3.5 levels. The preservation of small protrusion interdigitations in mutant lens fiber cells is likely facilitated by the remaining fimbrin, Arp3 and N-cadherin at the base of small protrusions (Cheng et al., 2016b).

### Levels of Tmod1, actin and some actin-interacting cytoskeletal components are reduced in Tpm3.5-deficient lenses

We examined whether changes in immunostaining of cytoskeletal proteins described above might be explained by changes in total or cytosolic vs. membrane-associated protein levels. We extracted proteins from pairs of 6-week-old *Tpm3/Δexon9d^+/+^* and *Tpm3/Δexon9d^−/−^* lenses and performed western blotting (Figure 10). Notably, there was a significant decrease in total Tmod1 levels (~50%), suggesting that reduced Tmod1 immunostaining signal on the fiber cell membrane (Figures 8-9) can be partly explained by a decrease in total Tmod1 protein levels. However, comparison of cytosolic and membrane-associated Tmod1 after biochemical fractionation of lenses showed that the proportion of Tmod1 associated with membranes was the same in *Tpm3/Δexon9d^+/+^* and *Tpm3/Δexon9d^−/−^* lenses (Figure 11). This suggests that Tmod1 puncta in the cytoplasm may be associated with transcellular F-actin populations that co-sediment with the lens membrane fraction, as discussed above.

**Figure 10.**
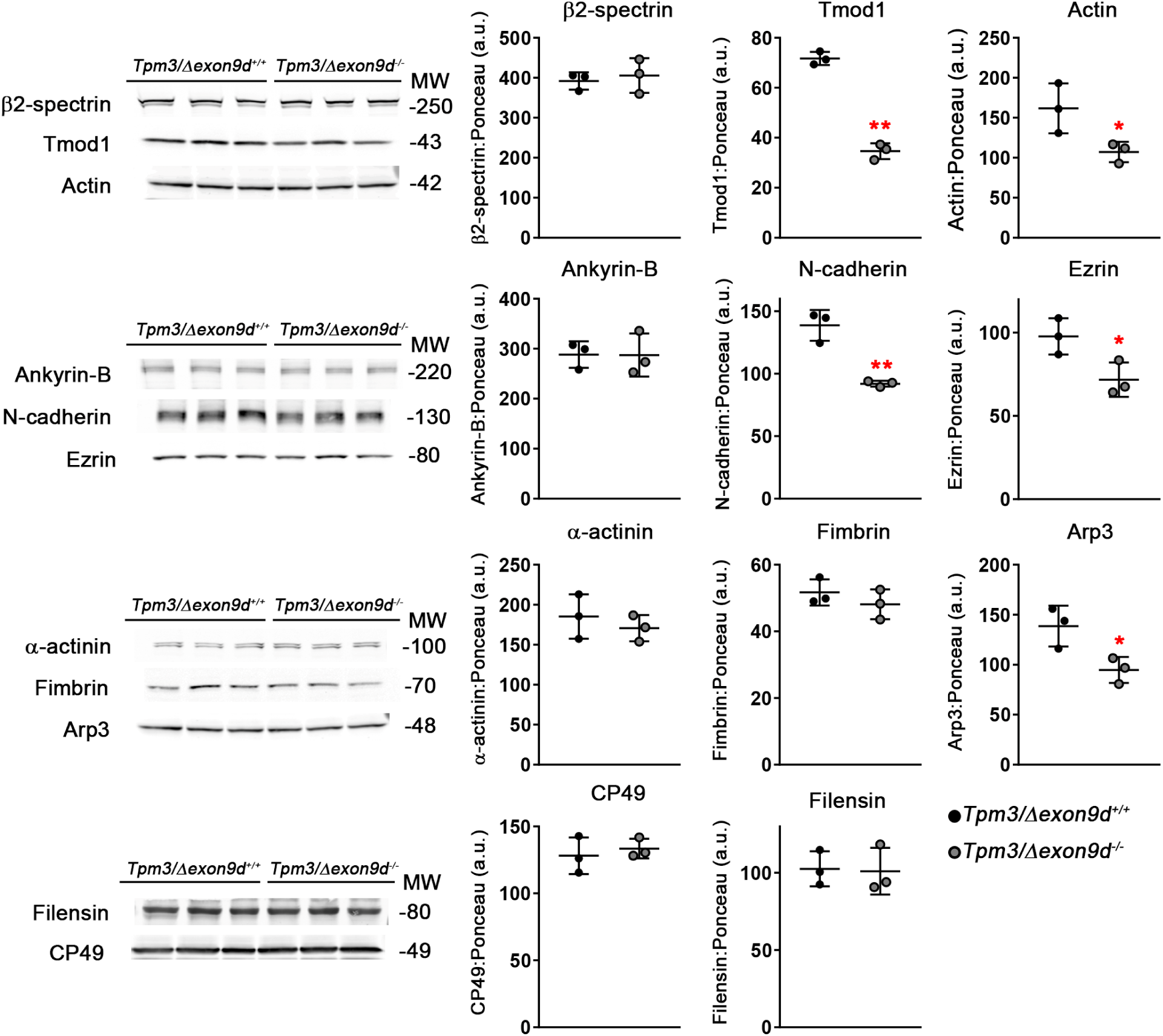
Reduced levels of Tpm3.5 affects levels of actin and select actin-associated proteins, Tmod1, N-cadherin, ezrin and Arp3. Western blots of indicated proteins from 6-week-old *Tpm3/Δexon9d^+/+^* and *Tpm3/Δexon9d^−/−^* whole lenses. All protein levels were normalized to total protein level (Ponceau S staining). Membrane skeleton components, Tmod1 and actin, are decreased in *Tpm3/Δexon9d^−/−^* lenses, while β2-spectrin levels remained unchanged. Actin-associated proteins that link F-actin to the cell membrane, N-cadherin and ezrin, have decreased levels in mutant lenses, while the level of ankyrin-B is unaffected. Arp3, a nucleating protein for branched F-actin networks, is decreased in mutant lenses, but levels of F-actin cross-linking proteins, u-actinin (Actnl) and fimbrin, are unchanged. Decreased Tpm3.5 does not affect beaded intermediate filament proteins, CP49 and filensin, levels. Plots reflect mean ± SD of n = 3 independent protein samples per genotype. *, *p*<0.05; **, *p*<0.01.

**Figure 11.**
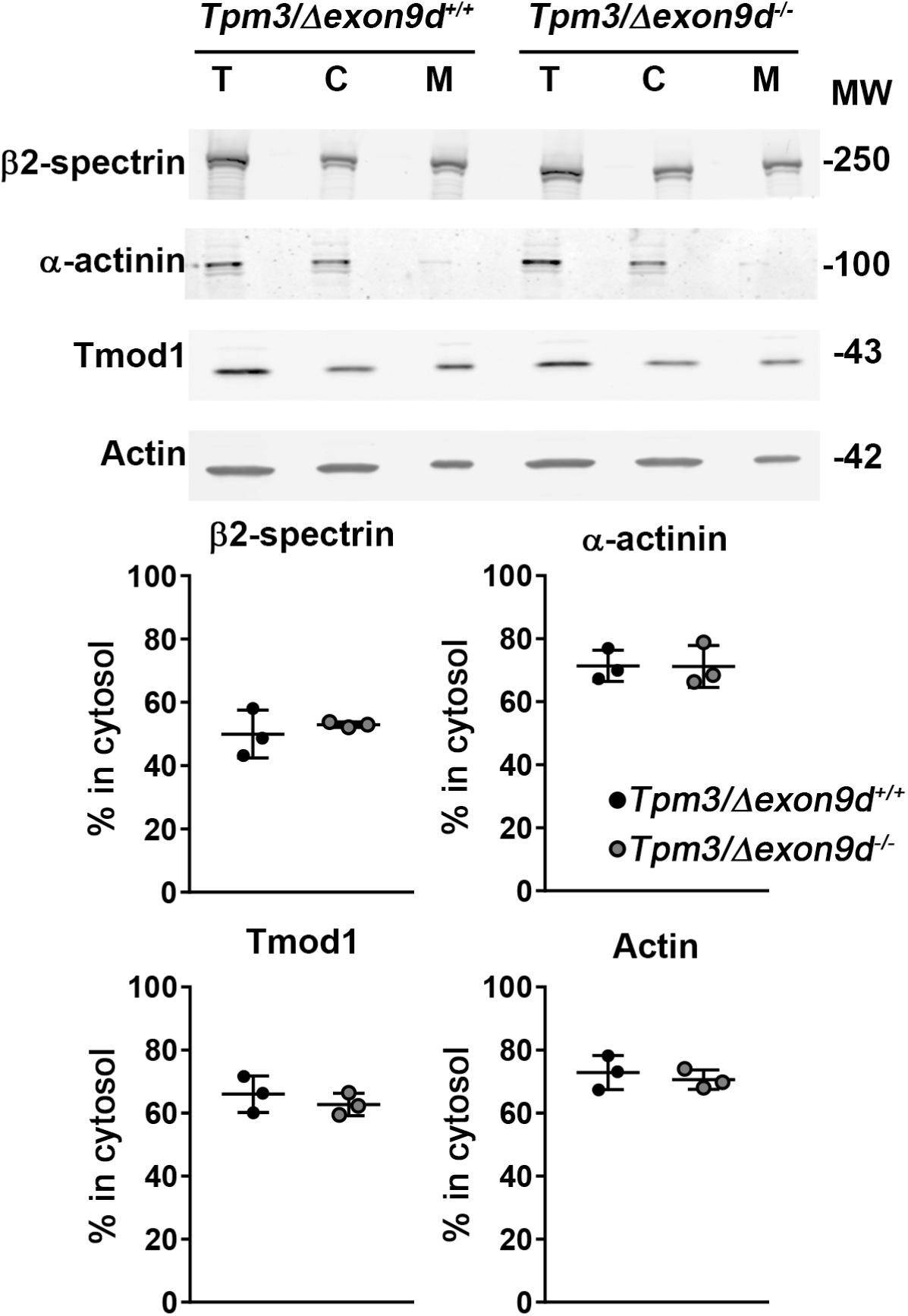
The ratio between cytosolic and membrane-associated fractions of actin and actin-binding proteins remains unchanged in *Tpm3/Δexon9d^−/−^* lenses. Western blots of total (T), cytosolic (C) and membrane (M) proteins from 6-week-old *Tpm3/Δexon9d^+/+^* and *Tpm3/Δexon9d^−/−^* lenses. The percent of each protein in the cytosol was calculated by dividing the cytosol band intensity by the total intensity of the cytosol plus membrane band. There is no change in the percent of actin, Tmod1, β2-spectrin and α-actinin (Actn1) in the cytosolic fraction of mutant lenses. Plots reflect mean ± SD of n = 3 independent protein samples per genotype.

We also observed a decrease in levels of total actin, N-cadherin, ezrin and Arp3 (Figure 10), which could suggest possible changes in cell-cell adhesion, although these decreases in total protein were not correlated with changes in the immunostaining signals nor in the subcellular distribution of any of these proteins (Figures 7–9 and S1). Conversely, striking changes in distribution of β2-spectrin and α-actinin (Actn1) in lens fiber cells observed by immunostaining are not correlated with changes in total protein levels. We also do not detect any change in the cytosolic vs. membrane-associated fractions of actin, β2-spectrin or α-actinin in mutant lenses (Figure 11). We also observed no changes in levels of ankyrin-B or fimbrin (Figure 10). However, the immunostaining result (Figure 9) suggests the fimbrin-associated F-actin network is partially disassembled. Finally, we checked whether disruptions of the actin cytoskeleton affect specialized lens beaded intermediate filaments, which are also important for lens mechanical properties (Fudge et al., 2011; Gokhin et al., 2012). There was no change in levels of CP49 or filensin that form beaded intermediate filaments (Figure 10). These data indicate that decreased Tpm3.5 leads to global changes in levels of actin and some but not all F-actin-associated proteins, which may weaken the actin cytoskeleton and contribute to reduced mechanical stability of lens fiber cells. Together, our immunostaining and Western blot results indicate that the F-actin network is rebalanced when Tpm3.5 levels are decreased, leading to a shift in the types of F-actin networks present in mutant lenses. While the proportion of F-actin associated with the fiber cell membrane remains the same, F-actin-cross-linking proteins appear to be redistributed to form alternate networks that do not contain Tpm3.5-coated F-actin.

## Discussion

We have demonstrated that Tpm is required for the normal biomechanical properties of the eye lens. This is the first evidence of the importance of Tpm for cell stiffness in a non-muscle mechanical-load-bearing tissue. Specifically, Tpm3.5, a previously unstudied isoform, is needed for normal lens stiffness and resilience. The function of Tpms in non-muscle cell biomechanical properties has been alluded to in studies of cultured cells and RBCs (Hundt et al., 2016; Jalilian et al., 2015; Sui et al., 2017; Tojkander et al., 2011; Wolfenson et al., 2016; Yang et al., 2016) but not previously shown *in vivo*. The lens is an ideal tissue to study the link between tissue biomechanics and cytoskeletal protein functions, because its function is tied to its mechanical properties. Other cytoskeletal components known to contribute to normal lens stiffness are the specialized beaded intermediate filament proteins, filensin and CP49, (Fudge et al., 2011; Gokhin et al., 2012), and the membrane skeleton proteins, periaxin and ankyrin-B (Maddala et al., 2016), but loss of these proteins also results in lens growth defects and/or fiber cell degeneration that complicates the interpretation of the mechanical property defects. In our case, the Tpm3.5-deficient lenses have a very mild shape change without any gross morphological defects, allowing a more straightforward understanding of the relationship between cytoskeletal structures and overall tissue mechanics.

### Decreased levels of Tpm3.5 lead to F-actin rearrangements, resulting in less mechanically stable networks and alterations in tissue mechanical properties

Our previous data and the work of others demonstrate that the actin cytoskeleton is vital to lens development, transparency, cell morphology, intercellular communication and biomechanics (reviewed by (Cheng et al., 2017). In *Tpm3/Δexon9d^−/−^* lenses, altered biomechanical properties are likely due to rearrangement of F-actin networks in lens fiber cells (Figure 12). Reduced Tpm3.5 is accompanied by dissociation of some Tmod1 from F-actin at the membrane, but an expansion of β2-spectrin- and α-actinin-associated F-actin networks along the cell membrane in mutant lens fiber cells. Partial dissociation of Tmod1 may be due to a decrease in F-actin-capping by Tmod1, since Tpms enhance Tmod1-F-actin-pointed-end capping affinity (Kostyukova and Hitchcock-DeGregori, 2004; Weber et al., 1994; Weber et al., 1999; Yamashiro et al., 2014). Expansion of both α-actinin-F-actin and β2-spectrin-F-actin networks could be a consequence of newly available F-actin binding sites that are normally occluded by Tpm3.5 (Christensen et al., 2017; Fowler and Bennett, 1984b). In addition, the cooperative assembly of loose F-actin networks may be formed by the homologous α-actinin, which contain spectrin repeats in its protein structure, and α2/β2-spectrin cross-linkers (Djinovic-Carugo et al., 2002; Winkelman et al., 2016). These expanded β2-spectrin and α-actinin networks without Tpm3.5-stablized or Tmod1-capped F-actin are likely less mechanically stable, leading to reduced *Tpm3/Δexon9d^−/−^* lens stiffness and resilience at high mechanical loads. This notion is consistent with *in vitro* studies showing that Tpms protect pointed-end depolymerization of F-actin (Broschat, 1990; Broschat et al., 1989; Weber et al., 1994) and that Tpm-coated F-actin is more mechanically rigid (Fujime and Ishiwata, 1971; Grazi et al., 2004; Kojima et al., 1994).

**Figure 12.**
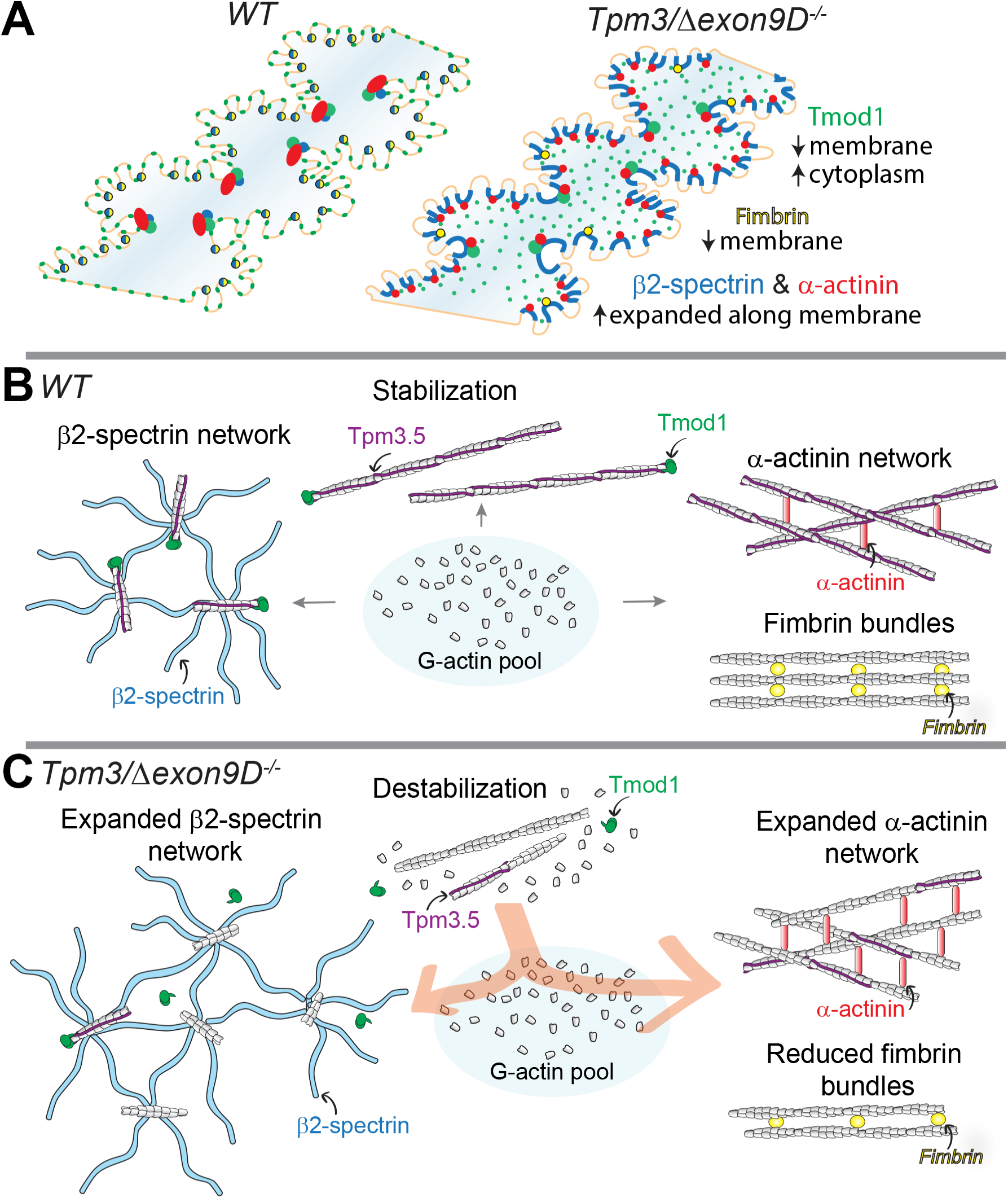
Tpm3.5 plays an important role in lens biomechanical properties likely by maintaining mechanically stable F-actin networks. **(A)** In *Tpm3/Δexon9d^−/−^* mature lens fibers with reduced Tpm3.5, Tmod1 (green) is dissociated from the cell membrane and appears as cytoplasmic puncta. Loss of Tpm3.5 and Tmod1 from F-actin at the membrane likely leads to expansion of the β2-spectrin (blue) and α-actinin (red) F-actin networks. Competition between fimbrin and α-actinin may also lead to the loss of fimbrin from F-actin at the membrane. **(B)** In normal fiber cells, there is a balance between Tpm3.5 and Tmod1 mechanically stabilized F-actin, the β2-spectrin-F-actin network and the α-actinin- or fimbrin-cross-linked F-actin networks. **(C)** In contrast, decreased levels of Tpm3.5 lead to depolymerization of Tmod1-capped F-actin. G-actin is then redistributed to other F-actin resulting in an expansion of the β2-spectrin and α-actinin networks. Not drawn to scale.

It is interesting to note that the loss of Tmod1 causes a different lens phenotype than Tpm3.5 deficiency. Tmod1 is required for the formation of complex interdigitations between mature fiber cells, and the loss of these paddles is correlated with a decrease in lens stiffness at low mechanical loads (Cheng et al., 2016b). In contrast, Tpm3.5-deficient lenses have normal fiber cell morphologies, but a decreased stiffness at high, but not low mechanical loads. In *Tmod1^−/−^* lenses, Tpm3.5 levels are reduced (Gokhin et al., 2012; Nowak et al., 2009), and the α-actinin-F-actin network is expanded (Cheng et al., 2016b), similar to the Tpm3.5-deficient lenses. However, β2-spectrin-F-actin networks at the membrane are disassembled in *Tmod1^−/−^* fiber cells (Cheng et al., 2016b), whereas they are expanded in Tpm3.5-deficient fibers. Since some Tmod1 remains associated with F-actin at the fiber cell membrane in Tpm3.5-deficient lenses (Figure 7), this may be sufficient to stabilize the β2-spectrin-F-actin networks at the membrane. Thus, F-actin networks in *Tmod1^−/−^* lenses (with reduced Tpm3.5 levels), and in Tpm3.5-deficient lenses (with reduced Tmod1 levels) appear to resemble one another in some, but not all respects, suggesting different functions for Tmod1 and Tpm3.5 in fiber cell F-actin networks, that then are reflected in their divergent functions in lens fiber cell morphology and biomechanics.

Our results are consistent with previous cell culture experiments where transfection with specific Tpm isoforms leads to assembly of different types of F-actin networks that alter cell migration and behavior, suggesting that Tpm isoforms control the type of F-actin networks that are formed (Bryce et al., 2003; Schevzov et al., 2005a; Tojkander et al., 2011). Importantly, the proportions of β2-spectrin, α-actinin, Tmod1 and actin associated with fiber cell membranes in biochemical assays remain unchanged in *Tpm3/Δexon9d^−/−^* lenses, suggesting that the ratio of cytosolic G-actin to polymerized F-actin and overall network assembly are not affected. Thus, reorganization of F-actin networks due to reduced Tpm3.5, does not appear to depend upon net F-actin polymerization or depolymerization. Our data, from an intact tissue, thus supports the prevailing hypothesis that the total pool of G-actin is stable and that a variety of F‑actin networks are engaged in competition for a limited pool of G-actin that is available for polymerization (Suarez and Kovar, 2016). The inter-network competition between F‑actin populations regulates network size and density, which can then determine cell and tissue biomechanical properties (Suarez and Kovar, 2016). While we have shown that decreased Tpm3.5 leads to steady-state F-actin network changes in fixed and immunostained lens fiber cells, it is unknown whether Tpm3.5 regulates active remodeling and reorganization of F-actin in lens fibers in resting lenses, or in lenses under load. Other studies have shown that Tpms can regulate the actomyosin contractile network and thus, influence cell mechanical properties and vesicular trafficking (Gunning et al., 2015; Khaitlina, 2015; Manstein and Mulvihill, 2016). Future studies to unravel interactions between Tpm3.5 and non-muscle myosins in the lens will be needed to determine whether Tpm3.5 regulation of the actomyosin network may contribute to lens biomechanical functions.

### Isoform compensation mechanisms in Tpm knockouts

Examination of Tpm isoforms expressed in the adult mouse lens by RT-PCR and sequencing of PCR products revealed expression of one long Tpm (Tpm1.7) and five short Tpms (Tpm1.8, Tpm1.9, Tpm1.13, Tpm3.5 and Tpm4.2) (Figures 1 and S2). Although Tpm3.1, which contains exon 9d, is not detected in the wild-type mouse lens, *Tpm3/Δexon9d^−/−^* lenses have an unexpected decrease in Tpm3.5, which contains exon 9a. This is unlike brain and red blood cells (RBCs) from *Tpm3/Δexon9d^−/−^* mice, which show expected reduction of Tpm3.1 (Figure 1 and (Fath et al., 2010; Sui et al., 2017)). Tpm3.5 mRNA and protein levels are both reduced in *Tpm3/Δexon9d^−/−^* lenses, suggesting deletion of exon 9d results in aberrant splicing of exon 9a. No compensation by increased expression of other Tpm isoforms from the Tpm3 gene, or from other Tpm genes, is detected in the mutant *Tpm3/Δexon9d^−/−^* lenses. However, it is certainly possible that normal levels of one or more of the other lens Tpms (1.7, 1.8, 1.9 or 4.2) may partially compensate for the reduced levels of Tpm3.5 in *Tpm3/Δexon9d^−/−^* lenses, attenuating the severity of observed lens defects. Future studies will be required to evaluate the functions of these other Tpms in the lens.

While an off-target isoform reduction has not been previously reported, several studies do show that isoform-specific knockout of Tpms can result in compensation by increased levels of a different Tpm isoform, likely accounting for mild phenotypes and complex disease mechanisms. Brain-specific knockout of exon 9c in *Tpm3* leads to compensation by exon 9a-containing isoforms, resulting in no change in the total level of Tpm3 proteins (Vrhovski et al., 2004), and no gross brain abnormalities. In *Tpm3/Δexon9d^−/−^* neurons and fibroblasts, loss of Tpm3.1/Tpm3.2 (exon 9d) is compensated by increased Tpm3 exon 9c isoforms, Tpm3.4 (aka TM5NM4) and Tpm3.7 (aka TM5NM7) (Fath et al., 2010). Only subtle changes in growth cone size and dendrite length were observed in cultured neurons isolated from *Tpm3/Δexon9d^−/−^* mice, suggesting that compensation by Tpm3.4 and Tpm 3.7 is sufficient to rescue the defects. In addition to alternative splicing compensation by products from the same gene, loss of one Tpm isoform can also result in functional compensation by another Tpm expressed from a different gene. RBCs only express Tpm1.9 (aka Tm5b) and Tpm3.1 (Fowler, 2013; Fowler and Bennett, 1984a; Sung et al., 2000; Sung and Lin, 1994), and loss of Tpm3.1 in *Tpm3/Δexon9d^−/−^* RBCs causes a compensatory upregulation of Tpm1.9, resulting in hyperstable F-actin and changes in the linkage between F-actin and the cell membrane (Sui et al., 2017). Therefore, tissue-specific compensatory upregulation or unexpected downregulation of alternate *Tpm* genes or splice products from the same gene can occur in exon-specific Tpm knockouts, and thus, knockout tissues may need to be screened more carefully for changes in other Tpm isoforms, looking for increased and decreased transcript and protein levels. More work needs to be done to unravel why germline deletion of exon 9d in *Tpm3* affects lens fiber cell transcript levels of a splice isoform that does not express the deleted exon.

### Decreased levels of Tpm3.5 results in a larger lens nucleus

This is the first report of a mutant mouse lens with an increased nucleus size, which is ~33% larger in volume in *Tpm3/Δexon9d^−/−^* lenses compared to controls. It has long been assumed that the hard lens nucleus affects overall lens tissue stiffness (Blankenship et al., 2007; Brown, 1973; Heys et al., 2004; Weeber et al., 2007). However, *Tpm3/Δexon9d^−/−^* lenses are softer but with a larger stiff lens nucleus, strongly suggesting the surprising conclusion that the lens nucleus does not play a significant role in the stiffness of the mouse lenses. This, in turn, implies that components determining the stiffness of mouse lenses under compressive load are restricted to the outermost fiber cell layers. This notion is supported by our recent study of multiscale load transfer in lens cells using a combination of coverslip compression of live mouse lenses and high resolution confocal microscopy. Under axial compression, we observe changes in the lens capsule, epithelial and fiber cell shape and fiber-fiber interactions at the periphery of mouse lenses (Parreno, et al. 2018, in press). More studies are required to reveal subtle changes in cellular size and organization during bulk lens shape change during equatorial stretch and to inform future mathematical models for lens shape change and stiffness.

The rigid lens nucleus is hypothesized to be formed by a complex and poorly understood process of cell compaction that involves remodeling of lens fiber cell membranes from large paddles and protrusions to smoother contoured membranes in inner fiber cells, with tongue-and-groove and square arrays of membrane-protein particles visible in freeze-fracture images (Al-Ghoul et al., 2001; Freel et al., 2003; Kuwabara, 1975; Lo and Harding, 1984). The change in fiber cell membrane architecture allows for compaction of inner fiber cells to form the hard nucleus (Costello et al., 2013; Taylor et al., 1996). However, we do not observe obvious changes in fiber cell membrane architecture by SEM between control and mutant lenses, suggesting that decreased Tpm3.5 levels do not accelerate the normal process of fiber cell membrane maturation and compaction. It is also hypothesized that inner fiber cell compaction is due to gradual loss of water from the cell cytoplasm driven by rearrangement of abundant crystallin proteins into larger aggregates, thus decreasing osmolarity (Kenworthy et al., 1994; Tardieu et al., 1992). This mechanism would imply that reduced levels of Tpm3.5 may affect ion and water homeostasis in the lens, perhaps via cytoskeletal regulation of Aqp0 or gap junction channel organizations. Interestingly, interactions between Tpm1.9 and aquaporin 2 water channels are needed for cell volume regulation in MDCK cells (Li et al., 2009). Further analyses of the *Tpm3/Δexon9d^−/−^* lens may provide a unique tool to elucidate how cytoskeletal proteins play a role in the innermost lens fiber cells during maturation and compaction to form the lens nucleus, as was originally proposed by Alcalá and Maisel (Alcalá and Maisel, 1985).

## Materials and methods

### Mice and lens pictures

All animal procedures were performed under an approved protocol from the Institutional Animal Care and Use Committee at The Scripps Research institute in accordance with the Guide for the Care and Use of Laboratory Animals by the National Institutes of Health.

*Tpm3/Δexon9d^−/−^* mice were a generous gift from Dr. Peter Gunning (University of New South Wales). Generation of *Tpm3/Δexon9d^−/−^* mice was previously described (Fath et al., 2010; Hook et al., 2011; Lees et al., 2013), resulting in an isoform specific knockout of exon 9d from *Tpm3*, resulting in absence of Tpm3.1, Tpm3.2 and Tpm3.13. Several mouse strains carry an endogenous mutation in the *Bfsp2/C*P49 gene that results in the loss of a specialized beaded intermediate filament in the lens (Alizadeh et al., 2004; Gokhin et al., 2012; Sandilands et al., 2004; Simirskii et al., 2006). Therefore, *Tpm3/Δexon9d^−/−^* knockout mice were backcrossed at least 12 generations to C57BL6/J wild-type mice that have wild-type *Bfsp2/CP49*. Heterozygous mice were bred to generate wild-type, heterozygous and homozygous knockout littermates, and genotyping for *Tpm3* and *Bfsp2/CP49* were performed by automated qPCR on tail snips (Transnetyx, Cordova, TN). Male and female wild-type and knockout littermates were used for experiments.

Six-week-old mouse lenses were dissected immediately from freshly enucleated eyeballs in 1X PBS (14190, Thermo Fisher Scientific, Grand Island, New York). Lens pictures were acquired with an Olympus SZ11 dissecting microscope using a digital camera.

### Antibodies and reagents

Unless otherwise noted, antibodies were used for both Westerns and immunostaining. All antibodies used in this study have been previously used and described. Unless otherwise stated, antibodies were diluted 1:100 or 1:1000 for immunostaining or Westerns, respectively. Rabbit polyclonal primary antibodies included anti-ankyrin-B (C-terminal), a generous gift from Dr. Vann Bennett (Duke University) (Scotland et al., 1998), anti-CP49 (rabbit 899) and anti-filensin (rabbit 76), generous gifts from Dr. Paul G. FitzGerald (University of California, Davis), anti-pan-fimbrin, a generous gift from Dr. Paul Matsudaira (National University of Singapore) (Correia et al., 1999), anti-GAPDH (sc-25778, 1:200 dilution for Westerns, Santa Cruz Biotechnology, Santa Cruz, California) and anti-human Tmod1 prepared in our laboratory (Gokhin et al., 2012). Mouse monoclonal primary antibodies used were anti-α-actinin (non-sarcomeric, Actn1, A5044, Sigma-Aldrich, St. Louis, Missouri), anti-actin (C4, 1:20,000 dilution for Westerns, Millipore, Burlington, Massachusetts), anti-Arp3 (A5979, Sigma-Aldrich), anti-ezrin (E8897, Sigma-Aldrich), anti-N-cadherin (for Westerns, 18-0224, Zymed, South San Francisco, CA), anti-β2-spectrin (612563, BD Biosciences, San Jose, California) and anti-tropomyosin exon 9a [CH1, detects Tpm3.5, developed by Jim Jung-Ching Lin and obtained from the Developmental Studies Hybridoma Bank, Iowa City, Iowa (Gokhin et al., 2012; Nowak et al., 2009)]. Rat monoclonal anti-N-cadherin antibody (1:20 dilution for immunostaining) was a generous gift from Dr. Dietmar Vestweber (Max-Planck-Institute for Molecular Biomedicine) (Dorner et al., 2005). Sheep polyclonal anti-tropomyosin 5NM1 and 5NM2 (detecting Tpm3.1 and Tpm3.2) was from Chemicon (AB5447, Temecula, CA).

For immunostaining, secondary antibodies were Alexa-488-conjugated goat-anti-rabbit (A11008, Thermo Fisher Scientific), Alexa-488-conjugated goat anti-mouse (115-545-166, minimal cross-reaction, Jackson ImmunoResearch, West Grove, Pennsylvania), Alexa-647-conjugated goat-anti-rat (112-605-167, minimal cross-reaction, Jackson ImmunoResearch) and Alexa-647-conjugated goat-anti-mouse IgG (A21236, Thermo Fisher Scientific). Rhodamine-phalloidin (R415, 1:50 dilution, Thermo Fisher Scientific) was used to stain F-actin, and Hoechst 33258 (B2883, 1:1,000 dilution, Sigma-Aldrich) stained nuclei.

For Western blots, secondary antibodies (1:20,000 dilution) were IRDye-680LT-conjugated goat anti-mouse (926-68020, LI-COR, Lincoln, Nebraska), IRDye-800CW-conjugated goat anti-rabbit (926-32211, LI-COR) and IRDye-800CW-conjugated donkey anti-goat (926-32214, LI-COR).

### RNA isolation and RT-PCR

RNA was isolated from 6-week-old mouse lenses, retina, brain, heart, skeletal muscle and kidney using Trizol (Thermo Fisher Scientific) according to manufacturer instructions. Two lenses or retinas from each mouse were pooled together into one RNA sample and homogenized in 100µl of Trizol for RNA isolation. Other tissues were cut into small chunks (~20-30mg) and homogenized in 0.5ml Trizol for RNA isolation. Reverse transcription was performed using the Superscript^TM^ III First-Strand Synthesis System for RT-PCR kit (Thermo Fisher Scientific) from equal amount of total RNA for each sample. PCR was performed with the same amount of cDNA from each sample in a 20µl volume reaction. PCR conditions for all primer sets were: denaturation at 95°C for 2 minutes, 40 cycles of denaturation at 95°C for 30 seconds, annealing at 54°C or 55°C (depending on the melting temperature of the primer set) for 30 seconds and elongation at 72°C for 1 minute, and a final 2 minutes elongation at 72°C. cDNA quality was verified by PCR for β-actin (forward: TGCGTGACATCAAAGAGAAG, reverse: GATGCCACAGGATTCCATA, product size ~200bp). Tpm primers were designed using the NCBI website (Table 1) to span entire Tpm genes. Equal volume of PCR product from each individual sample was loaded into 1% agarose gel for electrophoresis analysis. Bands were cut from the gel using clean razor blades, and PCR products were purified using the QIAquick gel extraction kit (Qiagen, Germantown, Maryland). Sanger sequencing was performed by Genewiz (La Jolla, California). RNA from tissues other than the lens were used as positive controls for Tpm PCR reactions as needed. For semi-quantitative PCR, cycles of denaturation, annealing and elongation were lowered to 30 for Tpm3.5 and 20 for G3DPH. G3DPH was a housekeeping control gene using previously described primers (Xia et al., 2010). Band intensities were quantified using ImageJ, and Tpm3.5 band intensity was background subtracted and normalized to G3DPH band intensity. Mean, standard deviation and statistical significance (Student T-test, two-tailed) were calculated using Excel and graphed using GraphPad Prism 7.

### Lens biomechanical testing and morphometrics

Stiffness and morphometrics of 8-week-old lenses were tested using sequential application of glass coverslips as previously described (Baradia et al., 2010; Cheng et al., 2016a; Gokhin et al., 2012). Briefly, freshly dissected lenses were transferred to a custom chamber filled with 1X PBS. Lenses were compressed by glass coverslips, and images of the uncompressed and compressed lens were taken under an Olympus SZ11 dissecting microscope with digital camera. After loading and unloading coverslips, the lens capsule was gently dissected away from the lens by fine forceps. Peripheral fiber cells were removed by rolling the lens between gloved fingertips. This removed the soft outer fiber cells and left a very hard, compact and round lens nucleus (center region of the lens). Images were taken of the lens nucleus for morphometric analysis. Image analysis was performed using ImageJ. To calculate strain, ε = (d-d_0_)/d_0_, where ε is strain, d is the axial or equatorial diameter at a given load, and d0 is the corresponding axial or equatorial diameter at zero load. Resilience was calculated as the ratio between the pre-compression axial diameter over the post-compression axial diameter. Lens volume was calculated as volume = 4/3×π×r_E_2×r_A_, where r_E_ is the equatorial radius and r_A_ is the axial radius. Lens aspect ratio was a ratio of the axial and equatorial diameters. Nuclear volume was calculated as volume = 4/3×π×r_N_3, where r_N_ is the radius of the lens nucleus. The nuclear fraction was calculated as a ratio between the nuclear volume and the lens volume. Strain curves and morphometrics were calculated in Excel and plotted in GraphPad Prism 7. Plots represent mean ± standard deviation. Student T-test (two-tailed) was used to determine statistical significance.

### Western blotting

Fresh lenses from 6-week-old mice were collected and stored at −80°C until homogenization. Two lenses from each mouse were pooled into one protein sample. At least three pairs of lenses of each genotype were used to make separate protein samples. Lenses were homogenized on ice in a glass Dounce homogenizer in 250µl of lens homogenization buffer [20 mM Tris, pH 7.4 at 4°C, 100mM NaCl, 1mM MgCl_2_, 2 mM EGTA and 10 mM NaF with 1 mM DTT, 1:100 Protease Inhibitor Cocktail (P8430, Sigma-Aldrich) and 1 tablet of PhosStop per 10ml buffer (04906845001, Roche) added on the day of the experiment] per 10mg of lens wet weight. Solubilized total lens proteins were transferred to a new Eppendorf tube. Half of the volume of total lens proteins were divided into a separate Eppendorf tube, and centrifuged at 21,130*g* for 20 minutes at 4°C. The supernatant, containing the cytosolic protein fraction, was collected into a new Eppendorf tube. Total and cytosolic protein samples were diluted 1:1 with 2X Laemmli sample buffer. The pellet, containing the membrane protein fraction, was washed twice with homogenization buffer and centrifuged at 21,130g for 10 minutes between washes. The pellet was solubilized in lens homogenization buffer diluted 1:1 with 2X Laemmli sample buffer. All samples were briefly sonicated and boiled for 5 minutes. Proteins were separated on 4–20% linear gradient SDS-PAGE mini-gels (Thermo Fisher Scientific) and transferred to nitrocellulose membranes. Gels were cut at the 150 kD marker. The bottom half of the gel was transferred in buffer (12.5mM Tris and 96mM glycine in ddH_2_O) with 20% methanol, and the top half of the gel was transferred in buffer with 0.01% SDS without methanol for 1 hour at 4°C. After transfer, nitrocellulose blots were incubated in 1X PBS for 1 hour at 65°C to increase binding of transferred proteins to the membrane. Membranes were then stained with Ponceau S (09189, Fluka BioChemica, Mexico City, Mexico), gently washed with ddH_2_O (until the protein bands are pink and the surrounding membrane is white) and scanned to reveal total protein levels in each lane. Blots were blocked for two hours at room temperature with 4% BSA in 1X PBS. Primary and secondary antibodies were diluted in Blitz buffer (4% BSA + 0.1% Triton X-100 in 1X PBS). Membranes were incubated in primary antibodies overnight at 4°C with gentle rocking and then washed with 1X PBS + 0.1% Triton X-100 (3 times, 5 minutes/wash) before incubation in secondary antibodies for 2 hours at room temperature in the dark with gentle rocking. After secondary antibody incubation, blots were washed again with 1X PBS + 0.1% Triton X-100 (4 times, 5 minutes/wash). Bands on blots were visualized on a LI-COR Odyssey infrared imaging system, and band intensities were quantified using ImageJ with background subtraction and then normalization to total proteins between 40-250 kDa (Ponceau S staining). For cytosolic and membrane fraction calculation, the cytosolic fraction band intensity was added to the membrane fraction intensity to equal the total intensity. The cytosolic fraction band intensity was then divided by the total intensity to determine the percent of proteins in the cytosolic fraction. Mean, standard deviation and statistical significance (Student T-test, two-tailed) were calculated using Excel and graphed using GraphPad Prism 7.

### Scanning electron microscopy (SEM)

SEM of mouse lenses were performed as previously described (Cheng et al., 2016b). Briefly, a small hole was made in the posterior of enucleated eyeballs from 2-month-old mice. Eyeballs were fixed in fresh 2.5% glutaraldehyde in 0.1M sodium cacodylate buffer, pH 7.3 at room temperature for 48-72 hours. Lenses were dissected from eyes, and each lens was fractured using a sharp syringe needle (26G) in the anterior-posterior orientation. This orientation exposes interlocking protrusions and paddles along the short sides of fiber cells. Lens halves were postfixed in 1% aqueous OsO_4_, dehydrated in graded ethanol, dried in a critical point dryer (Tousimis Inc., Rockville, MD), mounted on specimen stubs and coated with gold/palladium in a Hummer 6.2 sputter coater (Anatech Inc., Union City, CA). Images were taken with a JEOL 820 scanning electron microscope (JEOL, Tokyo, Japan). To ensure images were from comparable depths in different lenses, the center of the lens nucleus was used as a reference, and images were aligned based on measurements from the center outward to the lens equator. Four lenses from each genotype were examined, and representative images are shown.

### Immunostaining of frozen sections

Frozen sections from 6-week-old lenses were prepared as previously described (Cheng et al., 2016b). Briefly, a small opening was made at the corneal-scleral junction of freshly enucleated eyeballs to facilitate fixative penetration. Eyeballs were then fixed in freshly made 1% paraformaldehyde (15710, Electron Microscopy Sciences, Hatfield, Pennsylvania) in PBS at 4°C for 4 hours. Samples were then washed in PBS, cryoprotected in 30% sucrose and frozen in OCT medium (Sakura Finetek, Torrance, CA) in the cross-section orientation in blocks. Twelve-micron thick frozen sections were collected with a Leica CM1950 cryostat. Immunostaining of lens sections was conducted as previously described (Nowak et al., 2009; Nowak and Fowler, 2012). ProLong^®^ Gold antifade reagent (Thermo Fisher Scientific) was used to mount the slides. Confocal images were collected using a Zeiss LSM780 microscope (100X objective, NA 1.4, 1X zoom). The lens equator in sections was identified based on the thickness of the lens epithelium (Nowak et al., 2009). Staining was repeated on 3 samples from different mice for each genotype, and representative data are shown.

### Immunostaining of single lens fiber cells

Single lens fiber cell staining was performed as previously described (Cheng et al., 2016b) with some modifications. Lenses were dissected from eyes from 6-week-old mice. Fine forceps were used to carefully dissect away the lens capsule leaving the bulk mass of fiber cells. Decapsulated lenses were fixed overnight in 1% paraformaldehyde in PBS at 4°C. Lenses were then cut into quarters using a sharp scalpel along the anterior-posterior axis. Fine forceps were used to gently remove the hard lens nucleus from each lens quarter (~40% of the tissue). Lens quarters were post-fixed in 1% paraformaldehyde in PBS for 15 minutes at room temperature and then washed two times briefly with PBS. Next, lens quarters were permeablized and blocked using 3% normal goat serum, 3% bovine serum albumin and 0.3% triton X-100 in 1X PBS for 1 hour at room temperature. Lens quarters were immersed in primary antibody diluted with blocking solution, overnight at 4°C. Samples were washed 3 times, 5 minutes per wash, with 1X PBS + 0.1% Triton X-100 and incubated in secondary antibodies for 3 hours at room temperature. Finally, lens quarters were washed again with 1X PBS + 0.1% Triton X-100 (4 times, 5 minutes per wash). To maximize dissociation of single lens fibers, fine forceps were used to tease apart the lens quarters into small tissue chunks, and these lens fiber bundles were then mounted in ProLong^®^ Gold antifade reagent onto a glass slide with a 1.5 coverslip. Super-resolution confocal Z-stacks with 0.17μm steps were collected of single fiber cells using a Zeiss LSM880 confocal microscope with Airyscan (100X objective, NA 1.46, 2X zoom). Z-stacks were processed using the Airyscan Auto 3D method (strength 6.0) in Zen 2.3 SP1. Airyscan Z-stacks were further analyzed using Volocity 6.3, and noise reduction using the fine filter was applied to all channels. Fiber cell morphology was used to approximate the maturity of fiber cells. Staining was repeated on at least 3 lenses from 3 different mice for each genotype, and representative data are shown. All images presented are single optical sections through the cytoplasm of the fiber cell.

## Acknowledgments

This work was supported by grants from the National Eye Institute R21 EY 027389 (CC), R01 EY017724 (VMF and subcontract to W-KL) and R01 EY05314 (W-KL and subcontract to VMF). The authors declare no competing financial interests.

---

**Figure S1.**
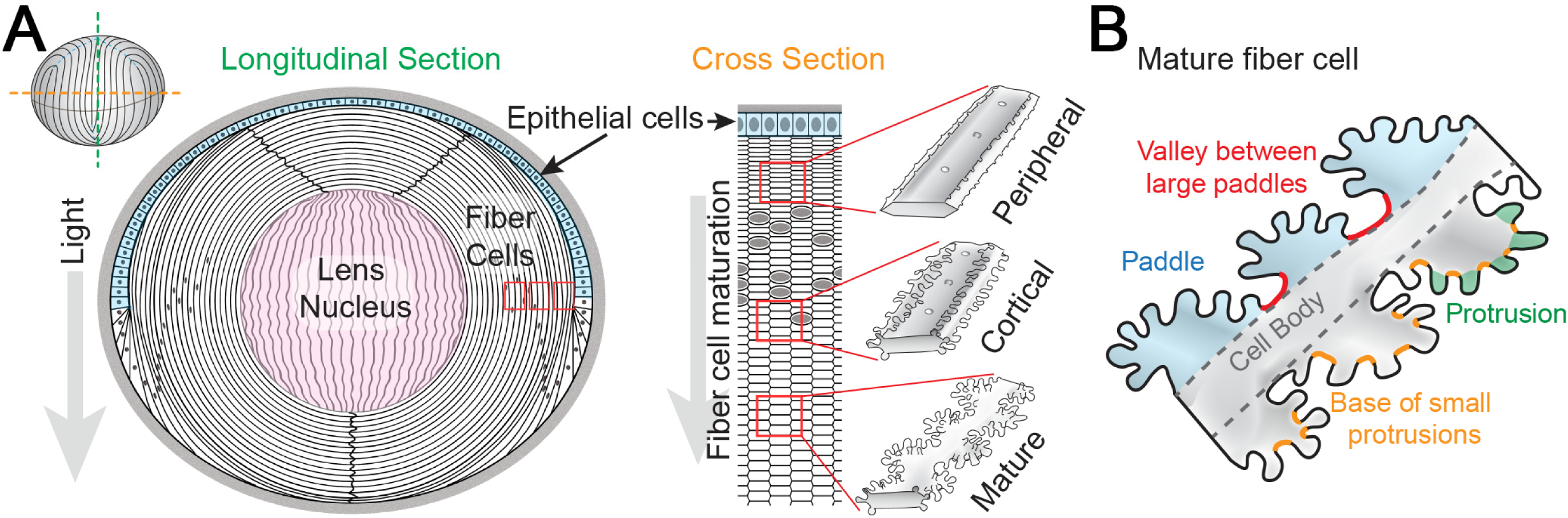
Lens anatomy diagrams. **(A)** A longitudinal section of the lens shows an anterior hemisphere of epithelial cells (blue) surrounding a bulk mass of lens fiber cells (white). The lens nucleus is composed of tightly compacted fiber cells in the middle of the lens (pink). A cross section through the lens reveals hexagon-shaped lens fibers packed into neat rows. As fiber cells start to differentiate, there are small protrusions along the short sides of these cells (peripheral cells). During maturation, cortical fiber cells form larger protrusions along the short sides. Mature fiber cells have large paddle domains decorated by small protrusions. These interlocking membrane interdigitations are thought to be important for mechanical stability in the lens. **(B)** A diagram of mature fiber cells with large paddle domains, the valley between large paddles, small protrusions and the base of small protrusions. Modified from (Cheng et al., 2016b). Not drawn to scale.

**Figure S2.**
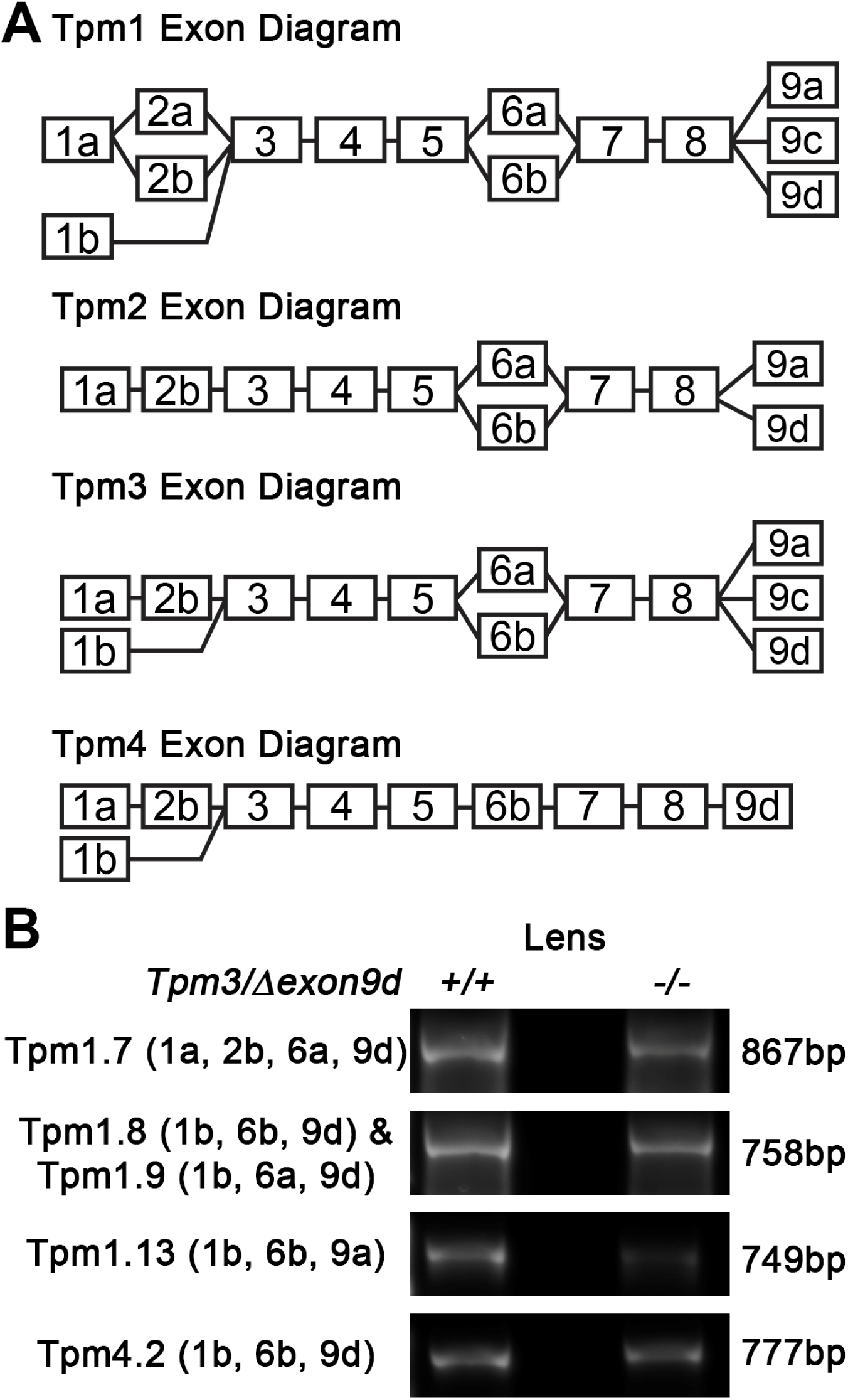
Five other Tpm isoforms (1.7, 1.8, 1.9, 1.13 and 4.2) are expressed at lower levels in the lens. **(A)** Diagrams of Tpm1-4 exons. Alternative splicing produces 20 known Tpm isoforms in mice. All Tpm isoforms contain exons 3, 4, 5, 7 and 8. Exons 1, 2, 6 and 9 differ between isoforms. Color codes correspond to Tpm isoforms found the lens in B. Not drawn to scale. **(B)** RT-PCR products for Tpms 1.7, 1.8, 1.9, 1.13 and 4.2 in 6-week-old *Tpm3/Δexon9d^+/+^* and *Tpm3/dexon9d^−/−^* lenses. These transcripts were expressed at lower levels than Tpm3.5, and upregulation of these isoforms in knockout lenses is not observed. No additional Tpms were detected in knockout lenses.

**Figure S3.**
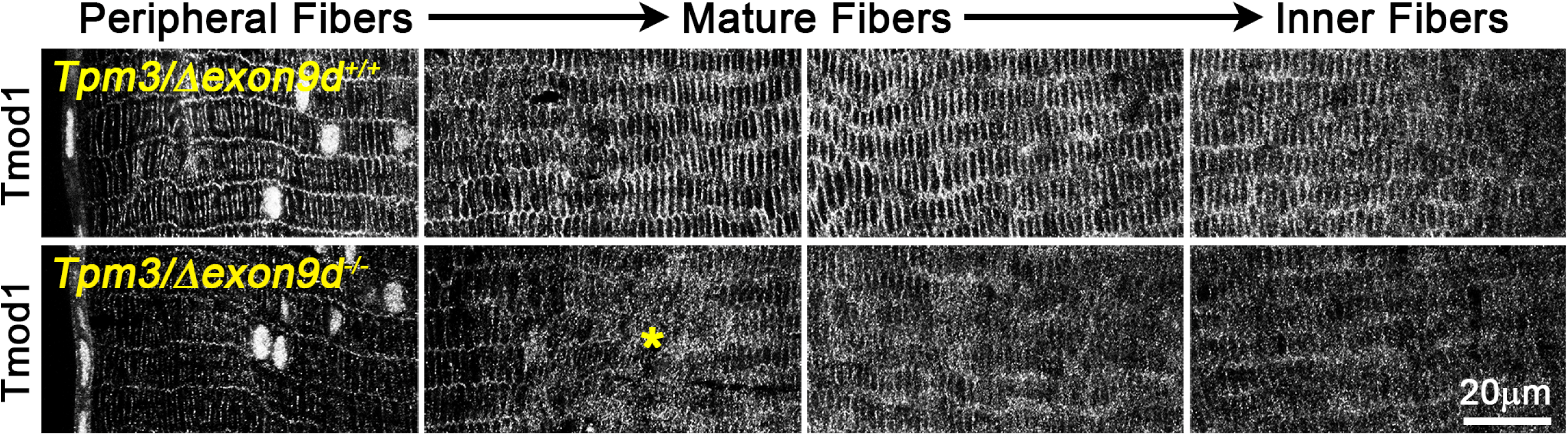
Tmod1 staining is disrupted in mature *Tpm3/Δexon9d^−/−^* lens fiber cells. Tmod1 immunostaining of frozen sections in the cross orientation from *Tpm3/Δexon9d^+/+^* and *Tpm3/Δexon9d^−/−^* lenses. Images are from sections near the lens equator from the lens periphery (leftmost) toward the inner fibers (rightmost). Tmod1 staining signal appears abnormally cytoplasmic in the mutant mature fiber cells (asterisk). Scale bar, 20µm.

**Figure S4.**
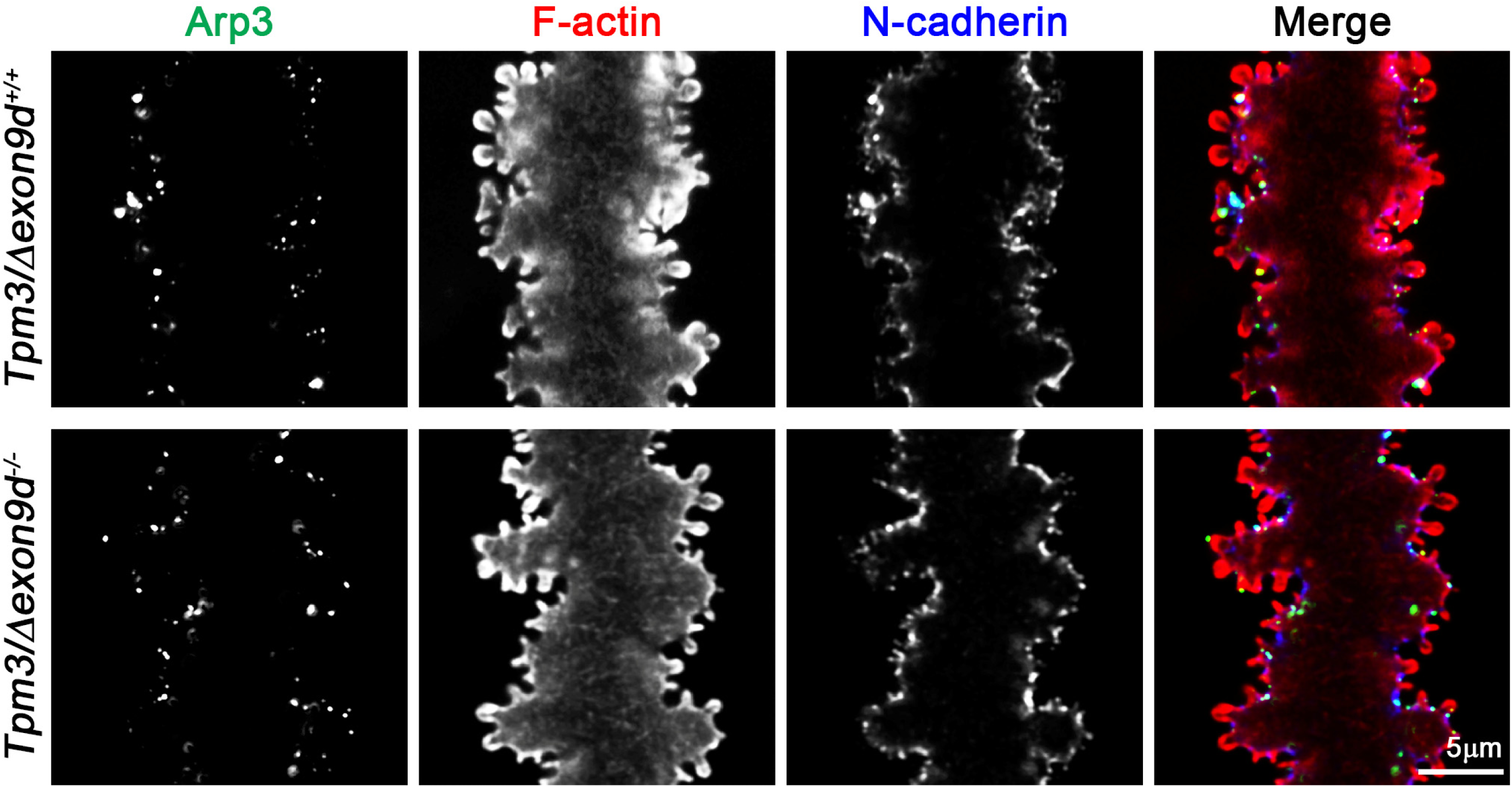
Arp3 and N-cadherin subcellular localization is not changed in *Tpm3/Δexon9d^−/−^* lens fibers. Immunostaining of single mature fiber cells from 6-week-old *Tpm3/Δexon9d^+/+^* and *Tpm3/Δexon9d^−/−^* lenses for Arp3 (green), F-actin (red) and N-cadherin (blue). Images are single optical sections through the cell cytoplasm. In control and mutant mature lens fibers, Arp3 and N-cadherin are localized in small puncta at the base of small protrusions. There is no obvious difference in staining pattern between control and mutant fiber cells.

